# A cellular and spatial atlas of *TP53*-associated tissue remodeling in lung adenocarcinoma

**DOI:** 10.1101/2023.06.28.546977

**Authors:** William Zhao, Benjamin Kepecs, Navin R. Mahadevan, Asa Segerstolpe, Jason L. Weirather, Naomi R. Besson, Bruno Giotti, Brian Y. Soong, Chendi Li, Sebastien Vigneau, Michal Slyper, Isaac Wakiro, Judit Jane-Valbuena, Orr Ashenberg, Asaf Rotem, Raphael Bueno, Orit Rozenblatt-Rosen, Kathleen Pfaff, Scott Rodig, Aaron N. Hata, Aviv Regev, Bruce E. Johnson, Alexander M. Tsankov

## Abstract

*TP53* is the most frequently mutated gene across many cancers and is associated with shorter survival in lung adenocarcinoma (LUAD). To define how *TP53* mutations affect the LUAD tumor microenvironment (TME), we constructed a multi-omic cellular and spatial tumor atlas of 23 treatment-naïve human lung tumors. We found that *TP53*-mutant (*TP53*^mut^) malignant cells lose alveolar identity and upregulate highly proliferative and entropic gene expression programs consistently across resectable LUAD patient tumors, genetically engineered mouse models, and cell lines harboring a wide spectrum of *TP53* mutations. We further identified a multicellular tumor niche composed of *SPP1^+^* macrophages and collagen-expressing fibroblasts that coincides with hypoxic, pro-metastatic expression programs in *TP53*^mut^ tumors. Spatially correlated angiostatic and immune checkpoint interactions, including *CD274*-*PDCD1* and *PVR*-*TIGIT*, are also enriched in *TP53*^mut^ LUAD tumors, which may influence response to checkpoint blockade therapy. Our methodology can be further applied to investigate mutation-specific TME changes in other cancers.

## INTRODUCTION

Non-small-cell lung cancer (NSCLC) is a heterogeneous disease that has routinely been classified into different histological subtypes^1^. More recently, genomic alterations have been used to characterize molecular subtypes of NSCLC that can be used to select treatments for patients with specific genomic changes^2–4^. Tyrosine kinase inhibitors (TKIs) and other targeted therapies against oncogenic mutations and chromosomal rearrangements (*e.g*., *EGFR*, *KRAS-G12C, ALK*) have led to a recent improvement in survival in NSCLC patients^5^. Treatments with immune checkpoint inhibitors (ICIs) (*e.g.*, anti-PD-1/PD-L1) with or without chemotherapy have proven efficacious in a subset of patients^6^ and are currently part of the first-line treatment in some patients with stage IV NSCLC^7^. However, due to high rates of cancer recurrence, more effective approaches toward patient stratification and precision therapy are urgently needed.

Tumor protein p53 (*TP53*) is the most commonly mutated gene in NSCLC (∼50%) as well as across many other cancers^8^. *TP53* mutations are more common in smokers^9^, and are associated with tumor progression and metastasis, leading to shorter survival in patients with NSCLC^10,11,12^. Despite being the most extensively studied tumor suppressor, *TP53* has been challenging to target therapeutically due to the diversity of mutations reported and its involvement in a wide range of regulatory pathways^13–15^. *TP53* is known to have an impact on the tumor microenvironment (TME) during tumor development in mouse models^16–18^. Mutant *TP53* is also associated with increased expression of PD-L1 and improved response to anti-PD1 therapy in human lung adenocarcinoma (LUAD)^19–24^, motivating us to investigate TME differences in LUAD patients with *TP53*-mutant (*TP53*^mut^) *vs. TP53*-wild type (*TP53*^WT^) tumors.

Single cell and spatial profiling of multiple treatment-naïve, resectable NSCLC patient tumors could advance our fundamental understanding of *TP53*-mutant cancer cells and their relationship to the TME and provide insights that may yield new therapeutic opportunities but, prior studies have not yet addressed this question. Single-cell RNA-sequencing (scRNA-seq) has previously been used to characterize malignant cells and the TME in resectable NSCLCs, focusing on gene expression differences between the primary tumor and adjacent normal tissue^25–28^, during tumor progression^29,30^, or across metastatic sites^31,32^. Studies have also leveraged TME characteristics to stratify NSCLC patients into clinically-relevant subgroups that could predict response to immunotherapy^22,33^. One study focused on characterizing the TME and intra-tumoral heterogeneity in *EGFR*^mut^ LUAD^34^, but these previous scRNA-seq studies did not determine the mutational status of *TP53* in tumors and did not investigate its impact on all cellular compartments in the TME.

Here, we generated the most comprehensive, multi-omic atlas of NSCLC to date, combining matched whole-exome sequencing, scRNA-seq, spatial transcriptomics, and multiplex immunofluorescence from *TP53*^mut^ and *TP53*^WT^ human tumors, to identify *TP53* mutation-associated changes in malignant cells and the TME. Our integrated atlas allowed us to identify spatially validated, cell-cell interactions that may contribute to shaping the tumor architecture of NSCLC. In doing so, we uncovered a hypoxic expression niche in *TP53*^mut^ LUAD and identified possible mechanisms by which malignant tumor cells may limit the growth of blood vessels in the TME. We further found a T cell compartment with highly exhausted features in *TP53*^mut^ *vs. TP53*^WT^ LUAD, which was accompanied by an enrichment in multiple immune checkpoint interactions between malignant, T, and myeloid cells. Finally, using a systematic approach to detect cellular landmarks in spatial transcriptomics data, we observed a potential pro-metastatic tissue niche consisting of *SPP1*-expressing tumor associated macrophages (TAMs) and a collagen-expressing population of cancer-associated fibroblasts (CAFs) linked to hypoxia in the *TP53*^mut^ LUAD TME. Overall, our approach provides a framework for leveraging multi-omic technologies to understand the impact of specific genomic alterations^35,36^ on the cellular and spatial architecture that can enable discovery of new biomarkers and more optimal methods for patient stratification in lung cancer and other tumor types.

## RESULTS

### A multi-omic atlas of NSCLC

To characterize the molecular, cellular, and spatial heterogeneity in NSCLC, we performed whole-exome sequencing (WES), scRNA-seq, spatial transcriptomics (ST; 10X Visium), and multiplex immunofluorescence (mIF) on primary tumor resections from 23 NSCLC patients (**Figure 1A**), of which 18 were diagnosed as LUAD (**Table S1**). Twenty-one of the 23 patients had a history of cigarette smoking (Median 40, Range 12-100 pack years) and all patients were confirmed to be treatment-naïve prior to tumor resection. We used WES analysis to determine the total tumor mutational burden, copy-number alterations (CNAs), and mutational status of genes commonly altered in NSCLC, including *TP53*, *EGFR*, *KRAS*, *STK11, KEAP1*, *RBM10*, and *PTPRD* in each tumor. There was high concordance between predicted CNA profiles from tumor-matched WES and those inferred from scRNA-seq data (**Figure 1B** and **Figure S1A**), many of which overlapped with commonly deleted (*e.g.*, *CDKN2A*, *PTPRD*, *B2M*) and amplified (*e.g.*, *NKX2-1*, *TERT*, *MCL1*, *EGFR*) genes in LUAD^8^. After filtering, we obtained a total of 167,193 high-quality cell profiles and used CNA inference to distinguish between malignant and non-malignant epithelial cells and then integrated^37^, clustered, and annotated them into 11 broad cell classes based on coherent expression of known marker genes (**Figure 1C**,**D**; **Figure S1B**,**C**; **Table S2**). Distinguishing tumors by their *TP53* mutation status, the different *TP53* mutations had comparable effects in downregulating the average expression of *TP53* target genes in malignant cells relative to *TP53*^WT^ LUAD tumors (**Figure 1E**). The remaining five tumors consisted of squamous, mucinous, and colloid histology, and were not used for comparisons between *TP53*^mut^ and *TP53*^WT^ tumors because *TP53* mutations have been found to have histology-dependent effects on patient outcome^38^.

**Figure 1.**
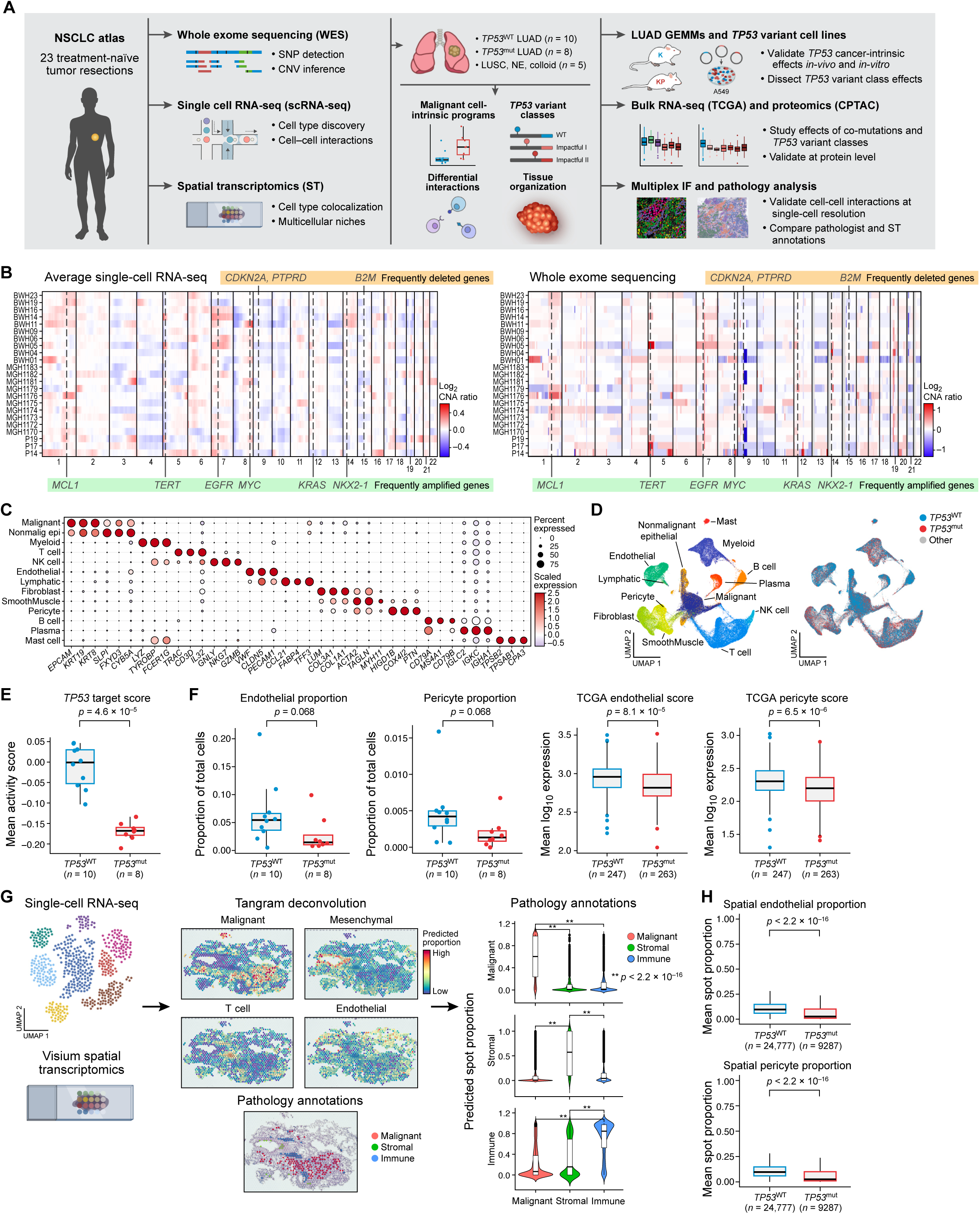
Spatially resolved, multi-omic atlas of NSCLC. **A.** Schematic of study design, including cohort selection (left), genomic assays profiled, sample stratification, downstream analyses, and validation platforms (right). **B.** Mean log_2_ copy number alteration (CNA) ratio inferred using scRNA-seq malignant cells (left) and bulk WES (right) from tumor-matched samples. Common focal amplifications (bottom, green) and deletions (top, orange) in LUAD are highlighted. **C.** Expression of representative markers across each annotated broad cell class. **D.** Uniform Manifold Approximation and Projection (UMAP) visualization of 167,193 cells, integrated by tumor sample to minimize intertumoral heterogeneity and colored by annotated cell classes (left) and *TP53* mutational status (right). **E.** Average *TP53* target gene expression score in malignant cells isolated from *TP53*^WT^ (blue) and *TP53*^mut^ (red) LUAD primary tumor samples. **F.** Left: Proportion of endothelial cells and pericytes from scRNA-seq of *TP53*^WT^ (blue) and *TP53*^mut^ (red) LUAD tumors. Right: Mean log_10_ expression of highly specific endothelial and pericyte markers derived from scRNA-seq in bulk RNA-seq data from TCGA. *P*-values were calculated using a two-tailed multiple regression t-test. **G.** Left: Integration of ST data with tumor-matched scRNA-seq data using Tangram allows for deconvolution of the proportion of different cell subsets. Middle: Spatial distribution of cell classes using Tangram (top), and a representative H&E stain annotated by a pathologist (bottom). Right: violin plots comparing proportion of computationally inferred cell classes (y-axis) within manually annotated categories (x-axis). Mann-Whitney-Wilcoxon test *p*-values were computed; Kruskal-Wallis FDR, *p* ≤ 0.05. **H.** Proportion of endothelial cells (top) and pericytes (bottom) in ST spots from *TP53*^WT^ (blue) and *TP53*^mut^ (red) LUAD sections. Mann-Whitney-Wilcoxon test *p*-values were computed.

### *TP53*^mut^ LUAD is associated with a distinct TME composition

Cell subsets in the TME of *TP53*^mut^ *vs*. *TP53*^WT^ LUAD tumors had distinguishable cell-intrinsic expression profiles (**Figure 1D**, **right**) and cell compositions (**Figure 1F** and **Figure S1D**). In particular, there was a pronounced decrease in the proportion of endothelial cells and pericytes in *TP53*^mut^ *vs*. *TP53*^WT^ LUAD tumors (**Figure 1F**, **left**), which we confirmed by scoring the corresponding cell type signatures in 510 bulk RNA-seq profiles of primary LUAD tumors from The Cancer Genome Atlas (TCGA) (**Figure 1F**, **right**). Endothelial cell abnormalities have been linked to tumor cell progression and metastasis^39^, and pericyte depletion has been linked to hypoxia-associated epithelial to mesenchymal transition (EMT) in breast cancer^40^.

These compositional differences between in *TP53*^mut^ and *TP53*^WT^ LUAD were also supported by spatial transcriptomic profiling of 14 tissue sections from 6 tumors in our cohort (2 with *TP53*^mut^ and 4 with *TP53*^WT^ LUAD). Spatial data from 2 additional tumors (6 slides) were excluded from downstream analysis due to either having an non-LUAD histology or an insufficient number of malignant cells (**Figure 1G**,**H** and **Table S1**). We used Tangram^41^, a deep learning framework, to map tumor-matched scRNA-seq profiles to the corresponding ST data and generate a probabilistic measure of the cellular composition for each spatial measurement (**Figure 1G**). Spatial cell composition inference was highly concordant with independent, manual annotations of the corresponding hematoxylin and eosin (H&E) stains by a pulmonary pathologist (N.R.M.; **Figure 1G** and **Figure S1E-G**). Consistent with our scRNA-seq, there was also a decrease in the proportion of endothelial cells and pericytes across ST spots in *TP53*^mut^ *vs*. *TP53*^WT^ tumor samples (**Figure 1H**).

### Dichotomous activity of malignant cell programs in LUAD associates with survival

While malignant cells from one tumor can occupy a gradient of expression states rather than discrete cell subsets^42,43^, different cancer expression programs and their usage across cells and tumors have not been comprehensively defined in LUAD at single-cell resolution. To identify shared cell states across tumors despite predominant, patient-specific differences in malignant cells^42,43^, we used canonical correlation analysis (CCA)^44^ to integrate and cluster malignant cells into 18 subsets shared across tumors (**Figure 2A** and **Figure S2A**) and annotated cell subsets using gene set enrichment analysis (**Figure S2B**), highlighting representative markers (**Figure 2B**). These subsets were enriched for genes associated with hallmark cancer processes (*e.g.*, cell cycle, hypoxia^45–47^, glycolysis^48^, partial EMT (pEMT)^49^, interferon gamma (IFNG) and TNFA signaling) and with lung epithelial cell identity (*e.g.*, alveolar type 2 (AT2-like), ciliated, secretory). Other malignant cell subsets showed marked expression of antigen presentation (MHCII), stress response^50^, metallothionein, and senescence-related genes^51^, which have not been previously identified *de novo* using scRNA-seq in NSCLC primary tumors. For downstream analyses, we defined LUAD malignant programs by the set of 10 most differentially expressed genes for each malignant subset and scored each program’s gene set across all malignant cells.

**Figure 2.**
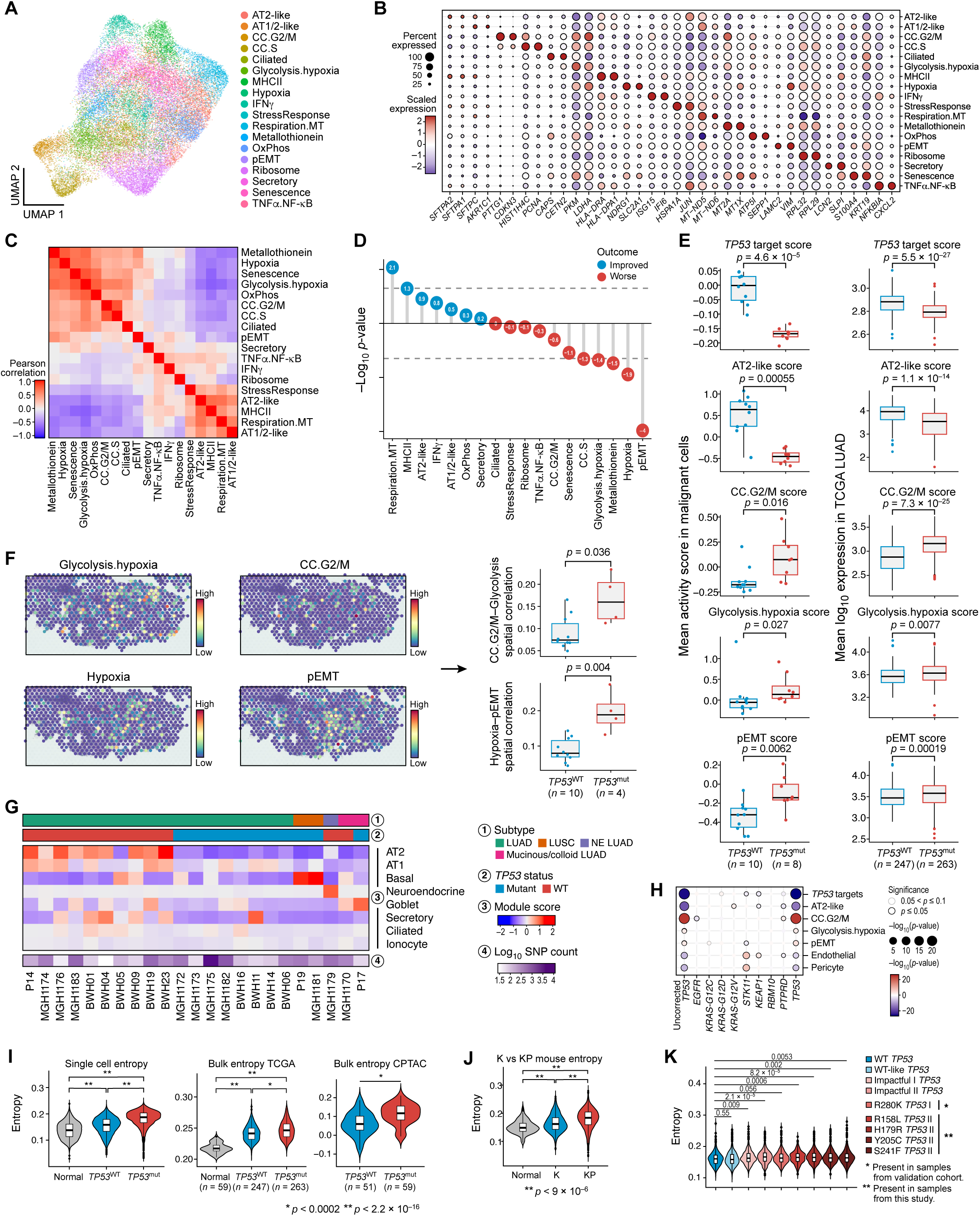
Enrichment of cell cycle, hypoxia and pEMT programs accompanied by loss of AT2 identity in *TP53*^mut^ LUAD malignant cells. **A.** UMAP of 33,377 malignant cells integrated across tumors and colored by annotated malignant expression program subsets. **B.** Dot plot of two selected markers representative of each malignant program subset. **C.** Pearson correlation of malignant program scores across all malignant cells. **D.** Lollipop plot showing -log_10_ *p*-values from Cox proportional-hazards regression analysis linking malignant programs to TCGA patient disease outcome, corrected for *TP53* mutational status. Sign of the y-axis corresponds to improved (positive value) or worse (negative value) outcome. **E.** Comparison of mean malignant program expression scores between *TP53*^mut^ and *TP53*^WT^ tumors in scRNA-seq (left) and in bulk RNA-seq data from TCGA (right). *P*-values were calculated using a Mann-Whitney-Wilcoxon test (scRNA-seq, left) or a two-tailed multiple regression t-test (TCGA, right). **F.** Left: Representative example of the spatial distribution of four malignant programs’ expression in a *TP53*^mut^ patient. Right: Bar plots comparing colocalization (spatial correlation) of pairs of malignant programs between *TP53*^mut^ and *TP53*^WT^ tumor sections. Mann-Whitney-Wilcoxon test *p*-values were computed. **G.** Module scores for normal lung epithelial cell signatures averaged across malignant cells in each tumor, ordered by histology (first) and *TP53* mutational status (second). Log_10_ SNP count for each tumor is shown on the bottom. **H.** Dot plot showing association of different mutations (columns) with mean log_10_ expression of malignant programs, endothelial, and pericyte scores (rows) in TCGA LUAD. Color and size of dots correspond to -log_10_ *p*-value assessed by Mann-Whitney-Wilcoxon (uncorrected *TP53*, left) or a two-tailed multiple regression t-test (*EGFR*, *KRAS-G12C*, *KRAS-G12D*, *KRAS-G12V*, *STK11*, *KEAP1*, *RBM10*, *PTPRD*, *TP53*). Black outline indicates *p*-value ≤ 0.05, and gray outline indicates *p*-value of ≤ 0.1. I. Distribution of entropy scores across non-malignant cells (grey), malignant *TP53*^WT^ cells (blue), and malignant *TP53*^mut^ cells (red) in the scRNA-seq data (left), across adjacent normal (grey), *TP53*^WT^ (blue), and *TP53*^mut^ (red) bulk transcriptomic LUAD tumor samples from TCGA (middle), and across *TP53*^WT^ (blue), and *TP53*^mut^ bulk proteomic tumor samples from CPTAC (right). *P*-values were calculated using a Mann-Whitney-Wilcoxon test. J. Distribution of entropy scores across normal cells (grey), malignant *Kras*^mut^*/ Trp53*^WT^ cells (K; blue), and malignant *Kras*^mut^*/Trp53*^mut^ cells (KP; red) in K and KP mouse tumors. K. Comparison of entropy scores between *TP53*^WT^ and different *TP53*^mut^ impact categories in A549 cells. Rightmost four violin plots show the entropy distributions of mutations also found in our cohort.

The programs clustered into two main groups after correlating their scores across all malignant cells (**Figure 2C**) and tumor samples (**Figure S2C**): one consisting of cell cycle, hypoxia, pEMT, and glycolysis programs, and another of alveolar-like, antigen-presentation, and stress response programs. Malignant program scores in these two groups were strongly anticorrelated, suggesting that they were regulated by two distinct and opposing pathways (**Figure 2C** and **Figure S2C**). This is consistent with a previous LUAD study where cells from patient-derived xenografts consisted of two distinct intra-tumoral subgroups, distinguished by a gene module consisting primarily of cell cycle genes^52^. High expression of several malignant programs (*e.g*., pEMT, hypoxia, cell-cycle, metallothionein, glycolysis) was associated with shorter survival across LUAD patient tumors in TCGA (**Figure 2D** and **Figure S2D**). In contrast, high expression of the antigen-presentation, MHCII program was linked to prolonged survival. Previously, various signatures derived from bulk RNA profiling in TCGA have been shown to be predictive of outcome in NSCLC^53,54^ but our *de novo* approach allowed for the discovery of more granular programs based on malignant cell expression only, unconfounded by other cells in the tumor microenvironment.

To identify cell-intrinsic differences between malignant cells from *TP53*^mut^ and *TP53*^WT^ LUAD, we compared the mean activity score of different malignant programs between the two patient groups and found higher expression in *TP53*^mut^ *vs*. *TP53*^WT^ LUAD for several programs associated with shorter survival (*e.g.*, cell cycle, glycolysis, pEMT), and a decreased expression of the AT2-like program and *TP53* targets (**Figure 2E**, **left**), consistent with the observation that *TP53*^mut^ tumors have shorter survival in LUAD TCGA (**Figure S2E**). There was not a statistically significant difference in either the number of single-nucleotide alterations (SNAs) or CNAs between *TP53*^mut^ and *TP53*^WT^ tumors in our cohort (**Figure S2F**). The malignant program differences were corroborated in bulk RNA-seq data from 263 *TP53*^mut^ and 247 *TP53*^WT^ LUAD primary tumors in TCGA (**Figure 2E**, **right**). These findings were also replicated when analyzing a compendium of independent cohorts (**Table S3**), assembled from published and unpublished scRNA-seq data from 22 additional treatment-naïve LUAD primary tumor samples with annotated mutational status (7 *TP53*^mut^ and 15 *TP53*^WT^)^26,32,55^ (**Figure S2G**). Notably, spatial co-expression of cell cycle with glycolysis and of hypoxia with pEMT programs was higher in *TP53*^mut^ *vs*. *TP53*^WT^ LUAD tumors (**Figure 2F**).

The decreased expression of the AT2-like program in *TP53*^mut^ LUAD was consistent with a loss of AT2 cell identity. Scoring each malignant cell profile across tumors for normal lung epithelial cell signatures^56^, *TP53*^WT^ LUAD malignant cells scored highly for AT2 markers (**Figure 2G**), whereas cells from tumors with squamous, neuroendocrine, or mucinous/colloid subtypes expressed the expected corresponding basal, neuroendocrine, or goblet signatures^57^. Interestingly, AT2 scores for *TP53*^mut^ LUAD malignant cells were significantly lower than *TP53*^WT^ LUAD but were comparable to AT2 scores in the tumors from non-adenocarcinoma subtypes, suggesting a loss of alveolar cell identity, which associates with tumor progression in LUAD^58^.

### Consistent cancer-intrinsic changes observed across different *TP53* variant and co-mutation classes

To understand the potential joint effect of *EGFR* and *KRAS* co-mutations on the observed *TP53*^mut^ LUAD cancer-intrinsic changes, we partitioned the TCGA cohort (n=506) into 6 subsets with distinct mutational statuses for these three genes (**Figure S3A**, **left**). *TP53*^mut^ tumors with or without co-mutations in *EGFR* or *KRAS* showed significant decreases in *TP53* target and AT2-like scores, and a significant increase in CC.G2/M score compared to *TP53*^WT^*KRAS*^WT^*EGFR*^WT^ tumors, whereas *TP53*^WT^ tumors with either *EGFR* or *KRAS* mutations did not have such significant differences in these signatures compared to tumors that were *TP53*^WT^*KRAS*^WT^*EGFR*^WT^. Interestingly, tumors with *TP53* mutations alone or in combination with *KRAS* but not with *EGFR* had significant increases in Glycolysis.hypoxia and pEMT scores compared to *TP53*^WT^*KRAS*^WT^*EGFR*^WT^ tumors, suggesting that *EGFR* mutations counteracted the *TP53*-induced effects on these malignant expression programs (**Figure S3A**), consistent with a recent report^59^. Multiple regression analysis also showed consistent cancer-intrinsic changes associated with *TP53* mutational status even when regressing out the effects of *EGFR*, *KRAS* (*G12C, G12D, G12V,* as well as all *KRAS* mutations together)*, STK11*, *KEAP1*, *RBM10*, and *PTPRD* mutations (**Figure 2H** and **Figure S3B**). Notably, there were no significant interaction effects between mutational status of *TP53* and *KRAS* or *TP53* and *EGFR* after ANOVA testing of model outputs.

To investigate the potential impact of different *TP53* mutational variants on changes to the expression of malignant gene programs, we partitioned the TCGA tumor samples based on dominant negative (DNE), loss of function (LOF), and Impactful *TP53* mutational classes as defined by previous functional studies overexpressing different *TP53*^mut^ variants in a *TP53*^WT^ A549 LUAD cell line^60,61^. Consistent with our tumor scRNA-seq analysis, decreases in *TP53* target, and AT2-like scores, and increases in CC.G2/M, Glycolysis.hypoxia, and pEMT scores were associated with TCGA LUAD tumors with different sub-categories of *TP53* DNE and LOF mutations (**Figure S3C**) or of *TP53* Impactful mutation sub-categories (**Figure S3D**). The DNE,LOF and Impactful II mutations gave rise to the most pronounced differences, where Glycolysis.hypoxia and pEMT programs are significantly increased only in DNE,LOF and Impactful II variant categories in TCGA (**Figure S3C** and **Figure S3D**). Moreover, many of these malignant cell intrinsic changes were also observed *in vitro* in A549 LUAD cells (*TP53*^WT^) overexpressing each of a broad spectrum of *TP53* mutational variants and profiled by Perturb-seq^61^ (**Figure S3E**). These trends held across all mutational categories, including all four *TP53* variants also found in our patients’ tumors and one variant also found in the scRNA-seq validation compendium (**Figure S3E**). As in TCGA, the malignant expression program changes were most pronounced for Impactful II mutations (**Figure S3E**) but were absent from A549 LUAD cells overexpressing different *KRAS* mutations (**Figure S3F**). Because we see consistent changes in malignant expression programs across a wide spectrum of *TP53* mutations, we reason that these phenotypic changes are associated with a loss of *TP53*^WT^ activity, rather than specific gain of function that arises from any specific mutation. In support of our observations, heterogeneous deletion events resulting in loss of *TP53* activity has recently been shown to result in homogenous and deterministic patterns genome evolution^62^.

Finally, we observed consistent changes in expression associated with *TP53* mutations at the protein level. Analyzing bulk proteomic data from the Clinical Proteomic Tumor Analysis Consortium (CPTAC) showed consistent decreases in *TP53* target and AT2-like signature scores, and an increase in CC.G2M signature scores at the protein level (**Figure S3G**), where effects were again most pronounced for DNE,LOF and Impactful II mutation categories (**Figure S3H**).

### Conserved changes in signaling entropy and malignant programs in *TP53*^mut^ human tumors and mouse models

We have previously reported loss of alveolar cell identity and reversion to a progenitor-like, highly-plastic cell state (HPCS) during LUAD progression in mice with somatic activation of oncogenic **<u>K</u>**RAS-G12D, which was greater in tumors from mice where the **<u>p</u>**53 was deleted (KP model) than those with wild-type p53 (K model)^63^. Consistent with our findings in human tumors, there was a significant decrease in AT2-like program score, and significant increases in CC.S, Hypoxia, Glycolysis.hypoxia, pEMT program scores when comparing KP versus K mice malignant cells (**Figure S4A**). Moreover, the expression of key hypoxia transcription factor *Hif1a* was higher in KP *vs*. K malignant cells (**Figure S4A**, **right**). Interestingly, we did not observe significant increases in the Hypoxia, Glycolysis.hypoxia, or pEMT programs between *TP53*^mut^ and *TP53*^WT^ A549 LUAD cells *in vitro* (**Figure S3E**), suggesting that the tumor microenvironment is important for shaping these malignant programs *in vivo*.

The consistent loss in alveolar type II cell identity in *TP53*^mut^ LUAD human tumors, cell lines, and mouse models prompted us to explore the consequences of this loss on signaling entropy^64^, a representative measure of cellular plasticity. We found an increase in signaling entropy in *TP53*^mut^ *vs. TP53*^WT^ LUAD malignant cells at the single-cell level in our cohort, which we confirmed in bulk RNA-seq and bulk proteomic profiles across a larger cohort size (**Figure 2I**), as well as in the KP *vs*. K mouse models^63^ (**Figure 2J**). This increase in entropy was most significant in DNE,LOF and Impactful II *TP53*^mut^ tumors in TCGA (**Figure S4B**) and was significant in *TP53*-only mutants and *TP53*-*KRAS* co-mutants but not in *TP53*-*EGFR* co-mutants in TCGA (**Figure S4C**), consistent with the trends observed in the malignant program activity changes.

Cell cycle and pEMT programs displayed the highest entropy amongst human malignant subsets (**Figure S4D**) and the greatest similarity to the HPCS cell state described in the KP mouse model (**Figure S4E**). Moreover, in A549 cells overexpressing mutant *TP53*, there was significantly increased entropy in cells harboring Impactful I and Impactful II *TP53* mutations, including variants found in our scRNA-seq and validation cohorts (**Figure 2K**). Conversely, there was no significant change in entropy in the same cells overexpressing different classes of *KRAS* mutants (**Figure S4F**), suggesting that the observed increase in entropy is directly linked to *TP53* mutations. Taken together, malignant cells from *TP53*^WT^ LUAD tumors were enriched for antigen presentation and AT2-like cell programs, while those from *TP53*^mut^ LUAD tumors were highly entropic, plastic, and associated with programs predictive of poor outcome, including cell cycle, glycolysis, hypoxia, and pEMT.

### *TP53*^mut^ LUAD is characterized by spatial interactions between malignant and stromal cells known to inhibit vascularization and promote EMT

We next focused on the decreased proportion of endothelial cells in *TP53*^mut^ LUAD (**Figure 1F**). This association is also observed in *TP53* mutant tumors in TCGA LUAD (**Figure 1F**), with or without co-mutations in *KRAS* and *EGFR* (**Figure S5A**, **left**), and is most significant in DNE,LOF *TP53* mutants (**Figure S5A**, **right**).

To investigate the potential mechanisms for decreased vascularization, we annotated 9 endothelial cell subsets based on the expression of established markers for endothelial cell subsets in the normal lung (**Figure 3A**, **Figure S5B-D**)^65^. These included: aerocytes, a specialized capillary cell responsible for gas exchange in the alveolus^66^ (expressing *HPGD*, *EDNRB*, and *SOSTDC1*); arterial cells (*GJA5*, *ENPP2*); capillary cells (*CA4*, *FCN3*); vascular endothelial cells (VEC), including *COL4A1*^+^ VECs (VEC.COL4A1), cycling (VEC.Cycling) and interferon-stimulated (VEC.IFN) subsets; lymphatic cells; pulmonary venous cells; and systemic venous cells. The proportion of aerocytes — and to a lesser degree arterial cells — out of all endothelial cells was decreased in *TP53*^mut^ *vs*. *TP53*^WT^ LUAD (**Figure 3B**), which was further supported by bulk RNA-seq data deconvolved using scRNA-seq-derived markers (**Figure S5E**). This suggests a corresponding decrease in gas exchange in *TP53*^mut^ LUAD and a resulting decrease in oxygen diffusion to the area directly surrounding the alveoli at the tumor site, providing a selective advantage for tumor cell survival^67^.

**Figure 3.**
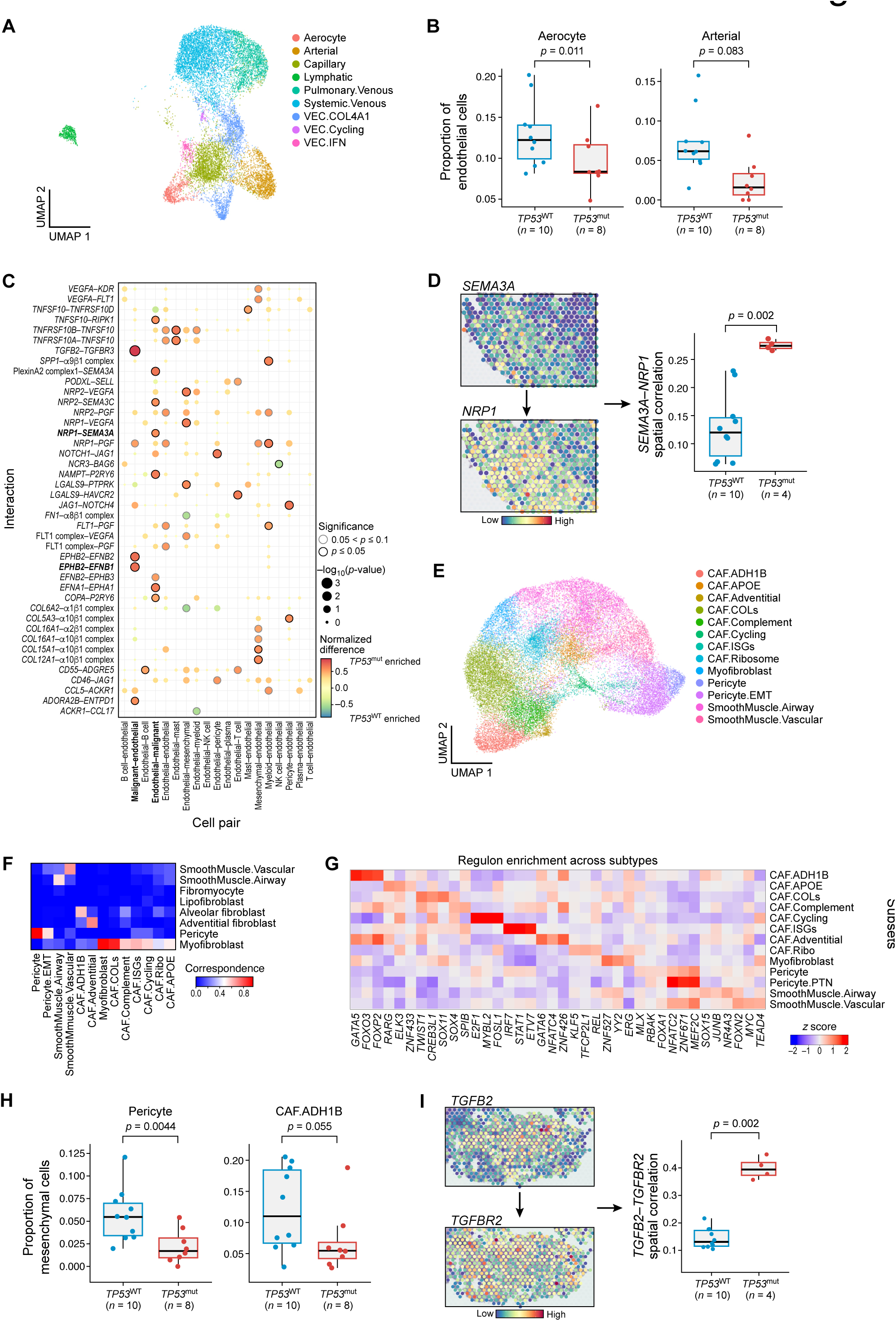
Spatially-resolved interactions linked to vascular depletion and stromal remodeling in *TP53*^mut^ LUAD. **A.** UMAP of 15,320 endothelial cells, integrated across tumors and colored by annotated subset. **B.** Comparison of aerocyte (left) and arterial (right) cell subset proportions in *TP53*^mut^ and *TP53*^WT^. Mann-Whitney-Wilcoxon test was used for computing *p*-values. **C.** Dot plot of differential ligand-receptor (rows) interactions between endothelial cells and other cell classes (columns). Red indicates enrichment in *TP53*^mut^ and blue enrichment in *TP53*^WT^ tumor samples. Black outline indicates *p*-value ≤ 0.05, as assessed by Fisher’s exact test. **D.** Left: Spatial expression of *SEMA3A* ligand and *NRP1* receptor in a representative *TP53*^mut^ ST sample. Right: Box plot comparing spatial correlation of *SEMA3A* and *NRP1* expression between *TP53*^WT^ and *TP53*^mut^ ST tumor samples. Mann-Whitney-Wilcoxon test *p*-value was computed. **E.** UMAP of 31,868 mesenchymal cells integrated across tumors and colored by annotated subset. **F.** Confusion matrix of correspondence between our *de novo* mesenchymal subset annotations and predicted identities using a classifier trained on annotated scRNA-seq data from normal lung mesenchymal cells. **G.** Z-scores of the most specific SCENIC regulon activity for each mesenchymal subset. **H.** Comparison of mesenchymal subset cell proportions in *TP53*^mut^ and *TP53*^WT^. Mann-Whitney-Wilcoxon test *p*-values were computed. **I.** Left: Spatial expression of *TGFB2* ligand and *TGFBR2* receptor in a representative *TP53*^mut^ ST sample. Right: Box plot comparing spatial correlation of *TGFB2* and *TGFBR2* expression between *TP53*^WT^ and *TP53*^mut^ ST tumor samples. Mann-Whitney-Wilcoxon test *p*-values were computed.

Analyzing ligand-receptor pairs, several interactions between malignant and endothelial cells known to inhibit endothelial cell growth and function were enriched in *TP53*^mut^ LUAD, including *SEMA3A*-*NRP1* and *EPHB2*-*EFNB1*^68–70^ (**Figure 3C** and **Figure S5F)**. Spatial transcriptomics from matched tumor samples support these observations, showing a significantly higher spatial correlation in the expression of *SEMA3A*-*NRP1* and *EPHB2*-*EFNB1* ligand-receptor pairs in *TP53*^mut^ *vs*. *TP53*^WT^ LUAD sections (**Figure 3D** and **Figure S5G**). Taken together, we observe spatially correlated expression of known inhibitory interactions enriched between malignant and endothelial cells in *TP53*^mut^ LUAD, which may explain the accompanying decrease in these subsets and vascularization.

The heterogeneity of mesenchymal cells and their potential role in faster tumor progression and shorter patient survival has been documented in NSCLC^26,71^. Mesenchymal cells in our tumors were partitioned into 13 subsets (**Figure 3E** and **Figure S6A-C**). Two subsets most closely resembled normal lung myofibroblasts^56^ (**Figure 3F**): CAF.COLs expressing high levels of collagens (*COL1A2*, *COL3A1*, *COL1A1*), which share markers with recently described *LRRC15*^+^ myofibroblasts^72^, and myofibroblasts, expressing both fibroblast and smooth muscle cell markers. We also annotated two pericyte-like subsets, one expressing canonical pericyte markers (*HIGD1B*, *COX4I2*), and one expressing *COL4A2* and EMT-related genes (**Figure S6A**,**B**); airway and vascular smooth muscle subsets, and alveolar (CAF.ADH1B) and adventitial (CAF.Adventitial) fibroblasts, which correspond to recently described human lung alveolar and adventitial fibroblasts^72^; and fibroblast subsets expressing higher levels of lipid metabolism (CAF.APOE), ribosomal (CAF.Ribo), complement and chemotaxis (CAF.Complement), and interferon-stimulated (CAF.ISGs) genes. Of note, we did not observe CAF subsets that resembled normal lung fibromyocytes or lipofibroblasts (**Figure 3F**). The different mesenchymal subsets were associated with different active regulons (**Figure 3G**), as inferred by pySCENIC^73^, with high regulon specificity (**Figure S6D**). As expected, cycling CAFs had an active *E2F1* regulon, a cell cycle regulator; CAF.ISGs activated the *IRF7* and *STAT1* regulons, controlling interferon responsive genes; and CAF.COLs activated a *TWIST1* regulon, previously linked to EMT in cancer^74^.

Based on the inferred composition of CAF subsets in spatial transcriptomics spots across sections, pericytes were significantly spatially correlated with endothelial cells (**Figure S6E**, Pearson correlation *P*-value < 10^-15^); CAF.COLs with vascular smooth muscle and CAF.ADH1B, CAF.Complement with CAF.Adventitial, pericytes with CAF.ISGs, and airway smooth muscle cells with CAF.APOE and CAF.Ribo. Spatially, most mesenchymal subsets were negatively correlated with malignant cells, indicating occlusion of fibroblasts from the tumor core (**Figure S6E**). Comparing the relative proportions of mesenchymal cell subsets (out of all mesenchymal cells) between *TP53*^mut^ *vs*. *TP53*^WT^ LUAD, pericytes and CAF.ADH1B were decreased in *TP53*^mut^ LUAD (as observed for broad cell classes) (**Figure 3H**), also supported by deconvolved bulk RNA-seq (**Figure 1F** and **Figure S6F**). Similar to endothelial scores, pericyte scores were significantly decreased in *TP53* mutant tumors in TCGA LUAD, with or without co-mutations in *KRAS* and *EGFR*, and this decrease was most significant in the DNE,LOF class of *TP53* mutations in TCGA (**Figure S6G**).

*TGFB2*-*TGFBR2* had significantly higher spatial correlation in *TP53*^mut^ *vs*. *TP53*^WT^ LUAD (**Figure 3I**), and *TGFB2*-*TGFBR2* and *TGFBR2*-*TGFB3* interactions between mesenchymal and malignant cells were enriched in *TP53*^mut^ *vs*. *TP53*^WT^ LUAD (**Figure S6H**), although the enrichment was not significant. *VEGFA* expression in myofibroblasts is higher in *TP53*^mut^ *vs*. *TP53*^WT^ LUAD (**Figure S6I**), consistent with previous reports that myofibroblasts can regulate angiogenesis in response to hypoxia^75^. Overall, changes in both stromal cell composition and spatial organization in *TP53*^mut^ LUAD are accompanied by an enrichment of putative interactions known to inhibit vasculature and induce epithelial-mesenchymal transition.

### Increased *SPP1* expression and immunomodulatory interactions in *TP53*^mut^ LUAD myeloid cells

Myeloid cells play an important role in tumor progression and may be a compelling therapeutic target in cancer^76^. While NSCLC myeloid cell populations were generally characterized^22,27^, the impact of *TP53* mutations on myeloid cells, and their interactions with malignant, stromal, and immune compartments in the TME has not been studied in detail in humans.

Myeloid cells partitioned into 14 subsets, largely consistent with previous NSCLC atlases^22^ (**Figure 4A,B** and **Figure S7A**), but with significant compositional differences between *TP53*^mut^ and *TP53*^WT^ LUAD. These include a significant decrease in *TP53*^mut^ LUAD in the proportion of TAM.FABP4 cells, which resemble tissue-resident alveolar macrophages (TAMs), and an increase in *SPP1*-expressing TAMs (TAM.SPP1) (**Figure 4C**). In agreement, *SPP1* expression was significantly increased both in TCGA bulk RNA-seq data and more prominently in myeloid pseudobulk comparisons between *TP53*^mut^ and *TP53*^WT^ LUAD tumors (**Figure 4D**, **left**). Expression of *CXCL9/10/11* was also increased in *TP53*^mut^ bulk TCGA LUAD tumors (**Figure S7B**). The proportion of *CXCL9/10/11* expressing TAMs (TAM.CXCLs) was also higher in *TP53*^mut^ LUAD tumors both in the scRNA-seq data and in deconvolution analysis of TCGA bulk RNA-seq data (**Figure S7C**). Notably, increases in *SPP1* and *CXCL9/10/11* were most significant in *TP53* mutant tumors without co-mutations in *EGFR* or *KRAS*, and in DNE,LOF and Impactful II classes of *TP53* mutant tumors in TCGA (**Figure S7D,E**).

**Figure 4.**
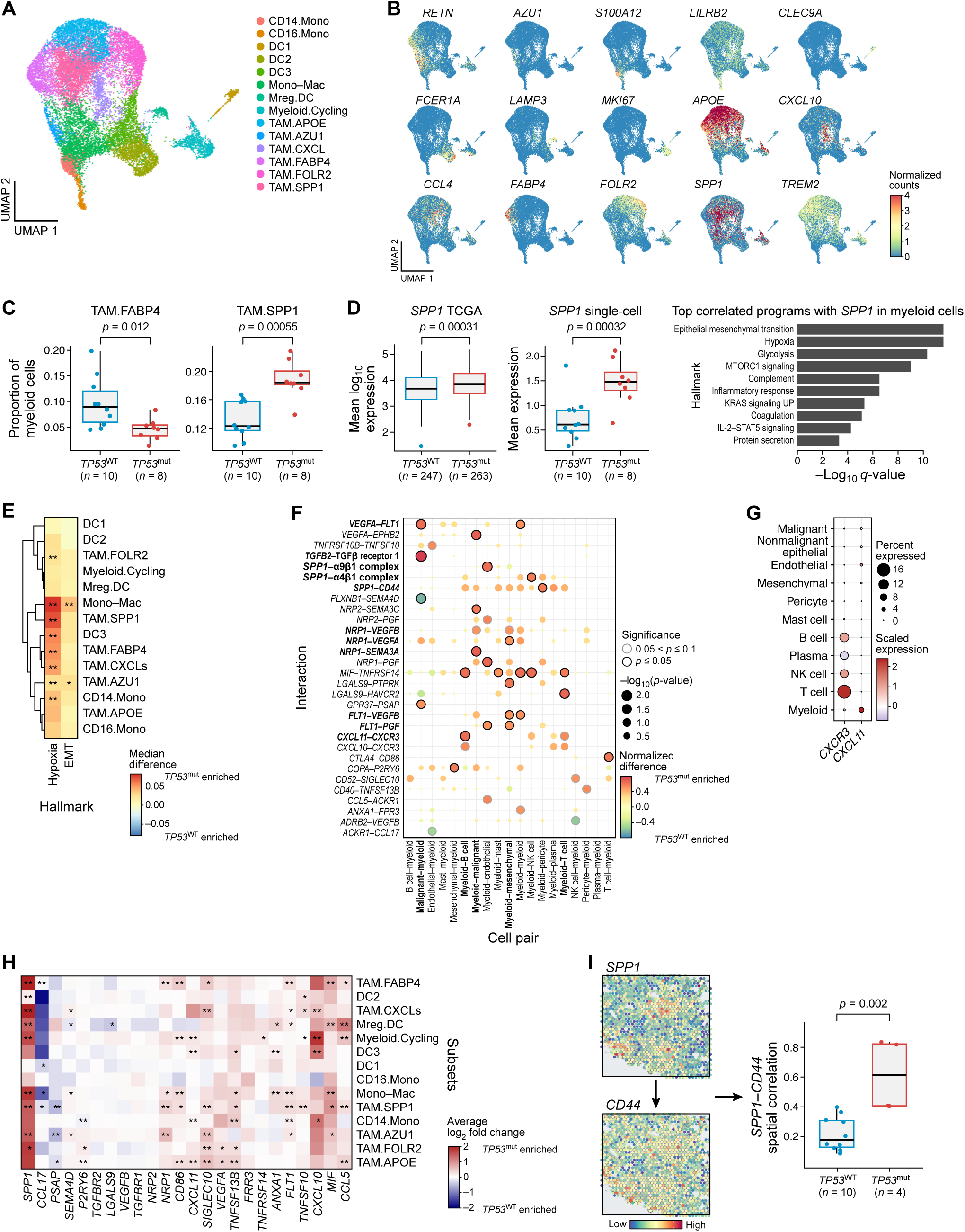
*SPP1* and other immunomodulatory factors are enriched in *TP53*^mut^ LUAD myeloid compartment. **A.** UMAP of 18,086 myeloid cells integrated across tumors and colored by annotated subset. **B.** Feature plot showing the expression of 15 representative myeloid subset markers. **C.** Comparison of myeloid subset cell proportions in *TP53*^mut^ and *TP53*^WT^ scRNA-seq tumors. Mann-Whitney-Wilcoxon test *p*-values were computed. **D.** Left: Comparison of *SPP1* expression in TCGA bulk RNA-seq (left) and scRNA-seq (middle) data from *TP53*^mut^ *vs*. *TP53*^WT^ primary tumor samples. Right: Horizontal bar plot of negative log_10_ *q*-values from gene set enrichment analysis of the 100 most positively correlated genes with *SPP1* in myeloid cells. *P*-values were calculated using a two-tailed multiple regression t-test (TCGA, left) or a Mann-Whitney-Wilcoxon test (scRNA-seq, middle). **E.** Differential enrichment of hallmark hypoxia and EMT module score for each myeloid subset, comparing pseudobulk subset profiles for *TP53*^mut^ *vs*. *TP53*^WT^ tumors. Mann-Whitney-Wilcoxon test *p-*values were calculated. **F.** Dot plot of differential ligand-receptor (rows) interactions between myeloid cells and other cell classes (columns). Red indicates enrichment in *TP53*^mut^ and blue enrichment in *TP53*^WT^ tumor samples. Black outline indicates *p*-value ≤ 0.05, as assessed by Fisher’s exact test. **G.** Dot plot of the expression of *CXCR3* and *CXCL11* across broad cell class annotations. **H.** Differential expression analysis of ligand and receptor genes from the interactions displayed in Figure 5F, comparing pseudobulk myeloid subset profiles for *TP53*^mut^ *vs*. *TP53*^WT^ tumors. Mann-Whitney-Wilcoxon test *p-*values were calculated. Red indicates enrichment in *TP53*^mut^ and blue enrichment in *TP53*^WT^ tumor samples. **I.** Left: Spatial expression of *SPP1* ligand and *CD44* receptor in a representative *TP53*^mut^ ST sample. Right: Box plot comparing spatial correlation of *SPP1* and *CD44* expression between *TP53*^WT^ and *TP53*^mut^ ST tumor samples. Mann-Whitney-Wilcoxon test *p*-values were computed.

Myeloid cell subsets in *TP53*^mut^ and *TP53*^WT^ LUAD were also associated with distinct cell-intrinsic gene programs, such that genes differentially expressed between myeloid subsets from *TP53*^mut^ *vs*. *TP53*^WT^ tumors were enriched for EMT, hypoxia, and interferon response pathway, especially in TAM.SPP1 cells (**Figure 4E** and **Figure S7F**). *SPP1* expression has been previously associated with EMT, and with early lymph node metastasis in LUAD^77^, as well as with shorter survival in TCGA LUAD independent of *TP53* mutational status (**Figure S7G**). Notably, the genes most correlated with *SPP1* expression across single myeloid cells were enriched for EMT, hypoxia and glycolysis functions (**Figure 4D**, **right**) and included VEGF receptors *NRP1* and *FLT1* (**Figure S7H**). In support of this observation, a recent compendium of 399 published scRNA-seq studies has shown that *SPP1*^+^ macrophages are consistently found in fibrotic lung diseases, COVID-19 lung, and pancreatic ductal adenocarcinoma (PDAC), suggesting a role of these fibrosis-associated *SPP1*^+^ macrophages in lung injury and ECM remodeling^78^. In summary, we find increased expression of *SPP1* in *TP53*^mut^ LUAD myeloid cells that is tightly linked with induction of genes involved in EMT and hypoxia programs.

To identify ligand-receptor interactions that may contribute to a distinct myeloid compartment in *TP53*^mut^ LUAD, we conducted differential ligand-receptor interaction analysis. Putative interactions enriched in *TP53*^mut^ LUAD include *VEGFA*-*FLT1* and *NRP1*-*SEMA3A* (**Figure 4F**), involved in recruitment of myeloid cells to tumor sites^79^, and *TGFB2*-*TGFBR1* interactions between malignant and myeloid cells, which are known to promote monocyte differentiation into TAMs^80^, and may help explain the relative increase in the monocyte-macrophage population in *TP53*^mut^ LUAD. Additionally, *NRP1*-*VEGFA*, *NRP1*-*VEGFB*, *FLT1*-*VEGFB*, and *FLT1*-*PGF* interactions between myeloid and mesenchymal cells, induced in *TP53*^mut^ LUAD, suggest that mesenchymal cells may contribute to monocyte recruitment to the TME (**Figure 4F**)^79^. In addition, *CXCL11*-*CXCR3* interactions between myeloid and B/T cells were enriched in *TP53*^mut^ LUAD (**Figure 4F**) with *CXCR3* specifically expressed by NK, B, and T cells (**Figure 4G**) and *CXCL11* induced in multiple myeloid subsets in *TP53*^mut^ LUAD (**Figure 4H**), most highly in TAM.CXCLs (**Figure S7I**), which constitute a higher proportion of myeloid cells in *TP53*^mut^ LUAD. Thus, myeloid cells may enhance recruitment of *CXCR3*-expressing T cells in *TP53*^mut^ LUAD. Finally, there was a significant increase in putative *SPP1*-mediated interactions between myeloid and endothelial cells or pericytes in *TP53*^mut^ LUAD (**Figure 4F**), also supported by increased spatial correlation of *SPP1* and *CD44* expression in *TP53*^mut^ tumors (**Figure 4I**). Thus, *TP53*^mut^ LUAD has increased expression of putative interactions involved in monocyte and lymphocyte recruitment, TAM differentiation, hypoxia, EMT, and other downstream effector functions *via SPP1, CXCL11*, and other TAM-associated immunomodulators.

### Enrichment of T cell exhaustion programs and putative immune checkpoint interactions in ***TP53*^mut^ LUAD**

Lymphoid cells in the scRNA-seq data partitioned into 13 subsets of T and NK cells (**Figure 5A,B** and **Figure S8A**), follicular and marginal zone B cells, IgA and IgG plasma cells, plasmacytoid DCs, and mast cells (**Figure S8B**). While there were no significant differences in B cell composition, *TP53*^mut^ LUAD had a decreased proportion of *CD4*^+^ tissue-resident memory T cells (CD4.TRM) and an increase in pre-exhausted-like^81^, *GZMK*-expressing *CD8*^+^ (CD8.GZMK) and exhausted-like (T.Exhausted) T cell populations out of all T and NK cells (**Figure 5C**), which was supported by our scRNA-seq validation compendium (**Figure S8C**) and by deconvolved bulk RNA-seq data from TCGA primary tumors (**Figure S8D**). Higher abundance of exhausted T cells may be mediated by increased chemotaxis *via CXCL11*-*CXCR3* interactions (**Figure 4F-H**), where *CXCR3* was upregulated in *TP53*^mut^ exhausted T cells (**Figure S8E**). We also observed a higher proportion of TFH cells relative to all T and NK cells in *TP53*^mut^ *vs. TP53*^WT^ tumors in both our discovery and validation scRNA-seq cohorts (**Figure S8C**), which was statistically significant when combining all samples. Expression of *CXCL13*, a top predictor for response to checkpoint inhibition therapy^82^, was most highly expressed in exhausted-like and TFH subsets and significantly increased in *TP53*^mut^ tumors (**Figure S8E**). Furthermore, expression of immune checkpoint molecules *PDCD1*, *CTLA4*, *HAVCR2*, and *TIGIT* were significantly higher across multiple T cell subsets in *TP53*^mut^ *vs. TP53*^WT^ tumors, while expression of *KLRB1*, a marker of favorable prognosis in cancer^83^, was downregulated (**Figure S8E**). Taken together, increased expression of *CXCL13* and immune checkpoint molecules (*PDCD1*, *CTLA4*, *HAVCR2*, and *TIGIT*) and proportions of TFH, *GZMK*-expressing *CD8*^+^, and exhausted-like T cells all suggest a heightened immunogenic potential in the *TP53*^mut^ LUAD TME that may contribute to a more favorable ICI immunotherapy response^82,84–86^.

**Figure 5.**
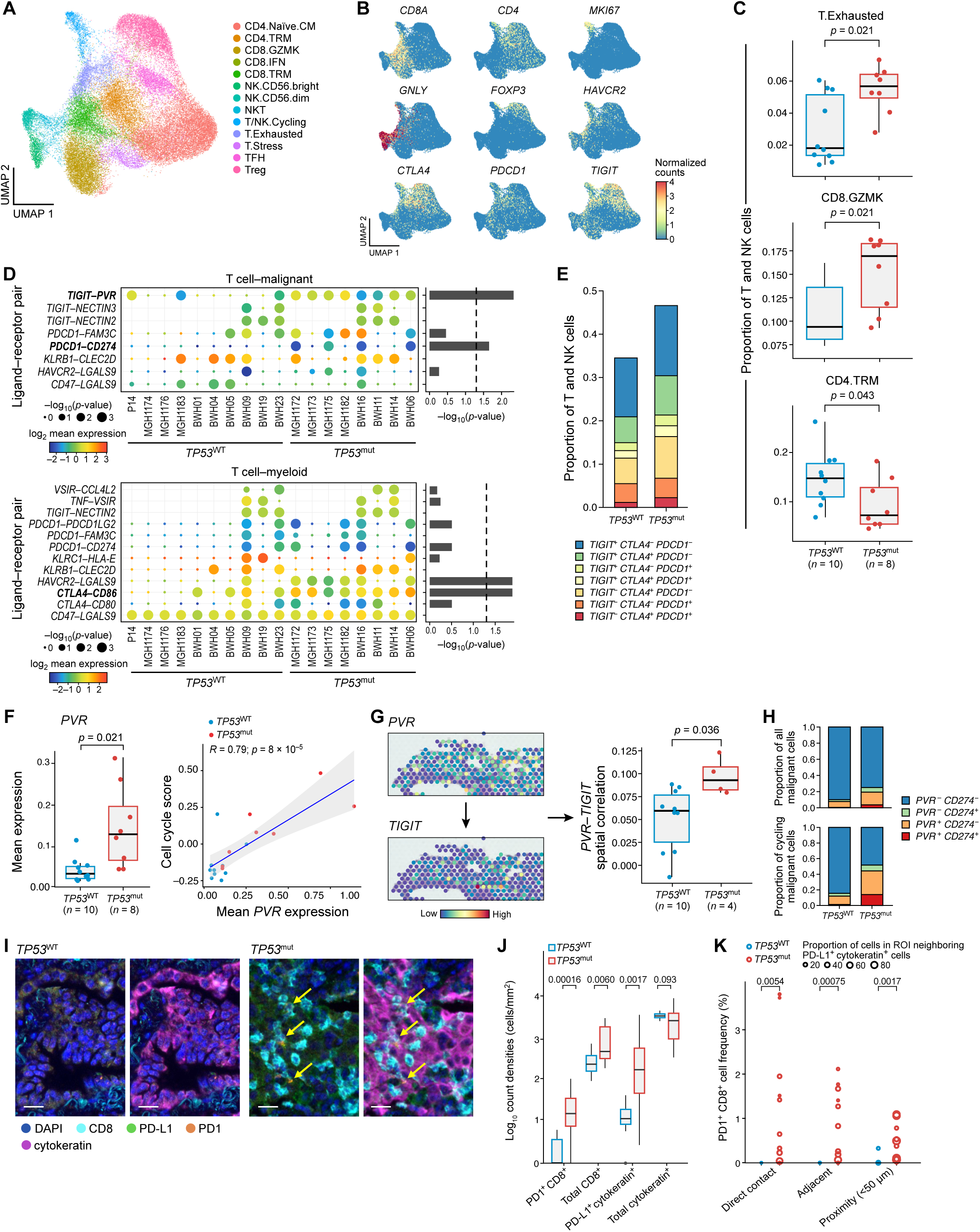
Immune checkpoint interactions and exhausted-like lymphoid cells are enriched in *TP53*^mut^ LUAD. **A.** UMAP of 40,293 T and NK cells integrated across tumors and colored by annotated subset. **B.** Feature plot showing the expression of 9 representative T and NK cell subset markers. **C.** Comparison of T.Exhausted, CD8.GZMK, and CD4.TRM T cell subset proportions relative to all T and NK cells in *TP53*^mut^ and *TP53*^WT^ scRNA-seq tumors. Mann-Whitney-Wilcoxon test *p*-values were computed. **D.** Log_2_ mean gene expression and predicted significance of ICI-targetable ligand-receptor interactions between T cells and malignant cells (top), and T cells and myeloid (bottom) for each tumor. Horizontal bar plots (right) show negative log_10_ *p*-value (Fisher’s exact test) of the enrichment in interactions between *TP53*^mut^ and *TP53*^WT^ tumors. **E.** Stacked bar plots depicting proportion of T and NK cells expressing different combinations of immune checkpoint molecules *TIGIT*, *CTLA4*, and *PDCD1* in *TP53*^WT^ (left) and *TP53*^mut^ (right) tumors. **F.** Left: Boxplot of mean *PVR* expression in *TP53*^WT^ (blue) and *TP53*^mut^ (red) tumor malignant cells. Right: Correlation of cell cycle score *vs*. mean *PVR* expression in malignant cells for each patient, where blue represents *TP53*^WT^ and red represents *TP53*^mut^ tumors. **G.** Left: Spatial expression of *PVR* and *TIGIT* in a representative *TP53*^mut^ ST sample. Right: Box plot comparing spatial correlation of *PVR* and *TIGIT* expression between *TP53*^WT^ and *TP53*^mut^ tumor ST samples (right) in the tumor periphery (spots containing ≤50% malignant cells). Mann-Whitney-Wilcoxon test *p*-values were computed. **H.** Stacked bar plots depicting proportion of malignant cells expressing different combination of *PVR* and *CD274* amongst all malignant cells (top) and amongst all cycling malignant cells (bottom) in *TP53*^WT^ (left) and *TP53*^mut^ (right) tumors. **I.** Representative mIF images of *TP53*^WT^ (left two; BWH01) and *TP53*^mut^ (right two; BWH06) tumors with and without cytokeratin (pink). Yellow arrows correspond to PD1^+^CD8^+^ cells. Scale bar, 20 µm. **J.** Box plots showing the log_10_ count density (cells/mm^2^) of PD1^+^CD8^+^, total CD8^+^, PD-L1^+^cytokeratin^+^, and total cytokeratin^+^ cells across regions of interest (ROIs) in *TP53*^WT^ (BWH01, BWH04, blue) and *TP53*^mut^ (BWH06, BWH11, red) tumor samples used for mIF. Mann-Whitney-Wilcoxon test *p*-values were computed. **K.** Grouped scatter plot showing the proportion of PD1^+^CD8^+^ cells out of all cells in proximity to PD-L1^+^cytokeratin^+^. Each circle represents an ROI from *TP53*^WT^ (BWH01, BWH04, blue) *vs*. *TP53*^mut^ (BWH06, BWH11, red) mIF tumor stains, where the size of the circle represents the proportion of PD-L1^+^cytokeratin^+^-neighboring cells, out of all cells within the ROI. Permutation Test *p*-values were computed.

Ligand-receptor analysis of putative immune checkpoint interactions^87^ between T, myeloid, and malignant cells showed an enrichment in *PDCD1*-*CD274* and *TIGIT*-*PVR* putative interactions between T and malignant cells in *TP53*^mut^ *vs*. *TP53*^WT^ LUAD (**Figure 5D**) and in *CTLA4*-*CD86* and *HAVCR2*-*LGALS9* putative interactions between T and myeloid cells. Indeed, we found higher expression of *CD274*, *CD86*, *PVR*, *PDCD1*, *CTLA4*, and *TIGIT* expression in *TP53*^mut^ bulk RNA-seq LUAD tumor samples from TCGA (**Figure S8F**), higher expression of *PVR* and *CD274* in *TP53*^mut^ bulk proteomic LUAD tumor samples from CPTAC (**Figure S8G**), and higher expression of *PDCD1*, *CTLA4*, and *TIGIT* in multiple T cell subsets in our scRNA-seq data (**Figure S8E**, **right**), where *TIGIT* was the most frequently expressed across T cells compared to *CTLA4* and *PDCD1* (**Figure 5E**). *PVR* expression was also higher in malignant cells from *TP53*^mut^ *vs*. *TP53*^WT^ tumor scRNA-seq data (**Figure 5F**, **left**) and highly correlated with the expression of the cell cycle program in malignant cells (**Figure 5F**, **right**), suggesting that *PVR* may be activated during cancer cell proliferation. Furthermore, there was significantly higher spatial colocalization of *TIGIT* and *PVR* expression in *TP53*^mut^ *vs*. *TP53*^WT^ LUAD (**Figure 5G**). In agreement, there was also a higher proportion of *PVR*^+^ malignant cells in *TP53*^mut^ *vs*. *TP53*^WT^ tumors (**Figure 5H**, **top**) and even more prominently in cycling malignant cells (**Figure 5H**, **bottom**).

To investigate the *TP53*^mut^ enrichment in interactions between PD-L1 (*CD274*) and PD1 (*PDCD1*) at a single-cell resolution, we applied multiplex immunofluorescence (mIF) to patient-matched *TP53*^mut^ and *TP53*^WT^ formalin-fixed paraffin-embedded (FFPE) slides from our cohort (**Figure 5I** and **Figure S8H**). There was a significant increase in log count density (cells/mm^2^) of PD-L1^+^cytokeratin^+^ cells, as well as PD1^+^CD8^+^ cells in *TP5*3^mut^ *vs*. *TP53*^WT^ tumor samples (**Figure 5J**), and, consistent with the scRNA-seq cell-cell interaction inference (**Figure 5D**), an increase in colocalization between PD-L1^+^cytokeratin^+^ cells and PD1^+^CD8^+^ cells in *TP53*^mut^ *vs*. *TP53*^WT^ tumor samples, across multiple spatial analyses measuring direct contact, adjacency (as defined by cells within the 6 nearest neighbors), and proximity (<50 μm) (**Figure 5K**). Taken together, increased expression of *PVR*, *TIGIT*, *CD274*, and *PDCD1* was concurrently observed with increased colocalization between the ligands and corresponding receptors in *TP53*^mut^ LUAD.

### Spatially-defined fibroblast-macrophage niche enriched in hypoxia and EMT programs in ***TP53*^mut^ LUAD**

To comprehensively map inter-compartment cellular colocalization using the spatial data, we deconvolved the proportion of cell subsets across all spatial transcriptomics spots and calculated the average correlation of cell type proportions across all LUAD sections and their difference between *TP53*^mut^ and *TP53*^WT^ LUAD tumors (**Figure 6A**, **Figure S9A**). Interestingly, there was a significantly lower colocalization (*i.e.,* more negative correlation) of endothelial cells with mesenchymal cells and with myeloid cells in *TP53*^mut^ *vs*. *TP53*^WT^ LUAD (**Figure 6B**), consistent with the increase in hypoxia related gene expression in mesenchymal and myeloid subsets in *TP53*^mut^ tumors. In addition, spatial colocalization of B, Plasma, and malignant cells was increased in *TP53*^mut^ LUAD (**Figure S9B**). B cell infiltration has been associated with increased PD1/PD-L1 expression, TMB, and prolonged survival in NSCLC^88^ and likely contributes to the heightened immunogenicity of *TP53*^mut^ LUAD tumors.

**Figure 6.**
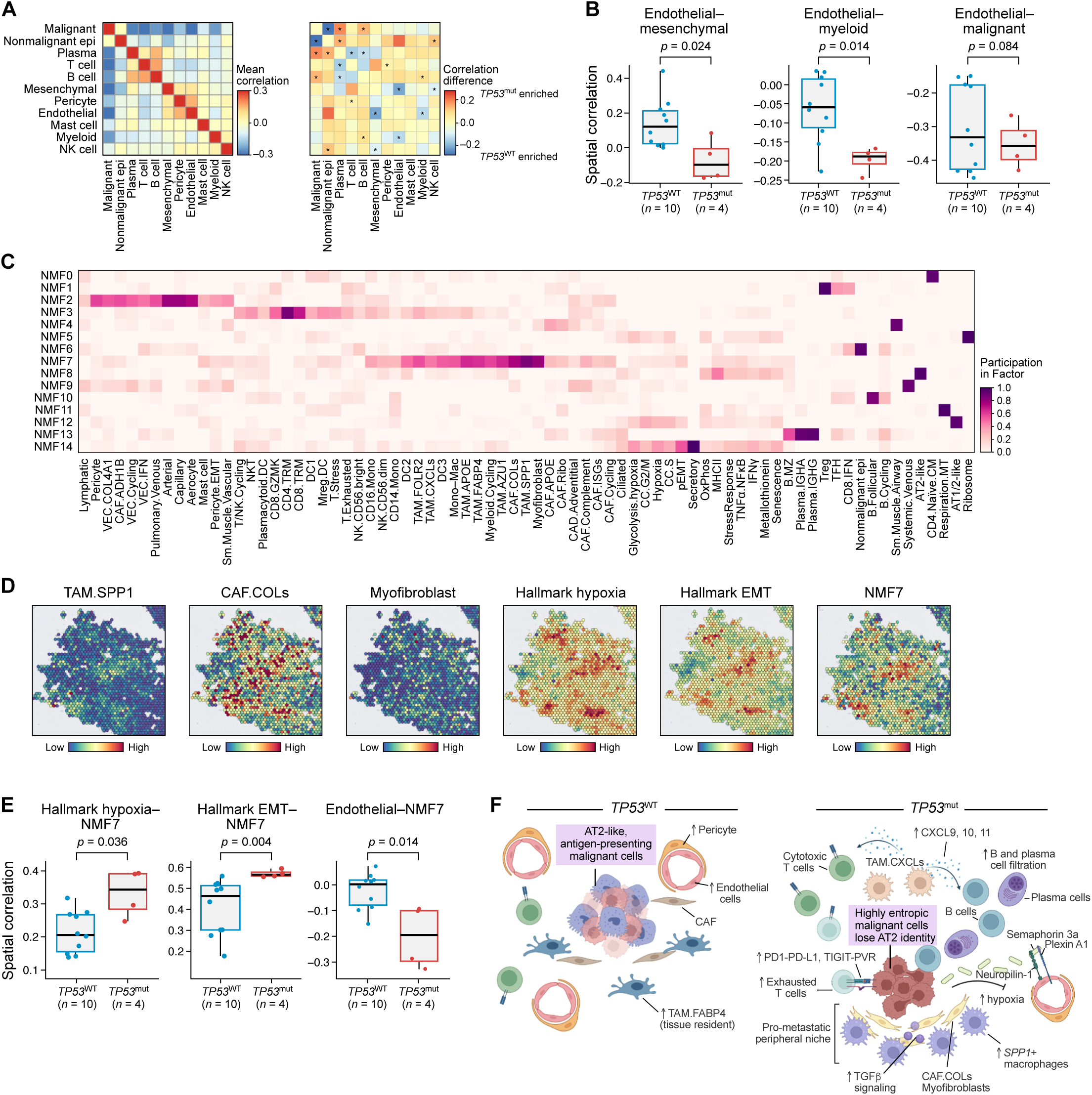
Spatial multicellular CAF-TAM niche enriched in hypoxia and EMT activity in *TP53*^mut^ LUAD. **A.** Overall spatial correlation of broad cell classes across ST spots (left) and differential spatial correlation between *TP53*^mut^ and *TP53*^WT^ tumor samples (right). Red indicates either high colocalization (left) or colocalization enrichment in *TP53*^mut^ (right), while blue represents either low colocalization (left) or colocalization enrichment in *TP53*^WT^ (right) tumor samples between pairs of cell subsets. **B.** Box plots comparing spatial correlation of endothelial cells with mesenchymal (left), myeloid (middle), and malignant (right) cells across spots in *TP53*^WT^ *vs*. *TP53*^mut^ ST tumor samples. **C.** Heatmap of the 15 factors depicting multicellular programs, as calculated using NMF. **D.** Spatial distribution of the enrichment of different myeloid and mesenchymal cell subsets comprising NMF7 (left) and of expression programs spatially correlated with NMF7 (right) in a representative *TP53*^mut^ tumor ST sample (patient BWH14). **E.** Box plots comparing spatial correlation of NMF7 with hallmark hypoxia score (left), hallmark EMT score (middle) and endothelial cell proportion (right) in *TP53*^WT^ *vs*. *TP53*^mut^ ST samples in the tumor periphery (spots with ≤50% malignant cells). **F.** Graphical summary of the differences between the *TP53*^WT^ (left) and *TP53*^mut^ (right) LUAD TME presented in this study.

We further identified multicellular communities by an NMF-based method to derive 15 factors with varying participation of different cell subsets (**Figure 6C**). Factor 7 (NMF7) contained multiple myeloid and mesenchymal subsets with known significance in tumor progression, including TAM.SPP1, CAF.COLs, and myofibroblasts, which were highly colocalized with each other, as well as with hallmark hypoxia and EMT program expression (**Figure 6D**). Spots high in NMF7 were low in malignant cell proportions and high in myeloid and mesenchymal cell proportions (**Figure S9C**). NMF7 was also more positively correlated with hallmark hypoxia and EMT program expression in the tumor periphery (spots with ≤50% malignant cells) and more negatively correlated with endothelial cells across all spots in *TP53*^mut^ *vs*. *TP53*^WT^ LUAD (**Figure 6E**). Because CAFs and macrophages are both suggested to induce EMT in tumors^89,90^, NMF7 may represent a hypoxic fibroblast-macrophage niche that resides on the periphery of the tumor core and may contribute to the early EMT phenotype in malignant cells in *TP53*^mut^ LUAD.

## DISCUSSION

Targeted therapies for NSCLC are currently directed at oncogenic driver mutations and chromosomal rearrangements (*e.g*., *EGFR*, *KRAS-G12C, ALK*), yet there are currently no effective therapies targeting *TP53* mutations, despite being common in tumors and extensively studied. Restoration of *Trp53* expression has been shown to cause tumor regression in mice^91^, leading to efforts to treat *TP53*^mut^ tumors by restoring wild-type p53 activity^14^. We hypothesized that cellular and spatial profiling of the TME in *TP53*^mut^ LUAD could provide a more detailed understanding of how *TP53* mutations could contribute to shorter survival and suggest new therapeutic opportunities. To this end, we characterized the LUAD TME of *TP53*^mut^ *vs*. *TP53*^WT^ tumors by combining scRNA-seq of 167,193 cells across 23 NSCLC patients, 23 matched tumor and normal WES, 20 spatial transcriptomics and 4 mIF tissue sections. Together, these represent the most comprehensive multi-omic map of NSCLC to date and provide a valuable resource of cell annotations, markers, and integrated data sets for the scientific community.

Our integrative analysis revealed a substantial remodeling of the TME associated with *TP53* mutations (**Figure 6F**), including decreased presence of pericytes and endothelial aerocytes. The mechanisms of TME remodeling are likely complex but our study uncovers multiple ones that may play a role. These include putative ligand-receptor interactions between malignant and endothelial cells (*SEMA3A*-*NRP1*, *EPHB2*-*EFNB1*) that were enriched in *TP53*^mut^ tumors and have known roles in inhibiting vascularization, and which were significantly more colocalized in the *TP53*^mut^ spatial data. *TP53*^mut^-specific depletion of endothelial cells was accompanied by increased hypoxia and pEMT program expression in malignant cells, which was altered across multiple functional categories of *TP53* mutations in bulk transcriptomic and proteomic data, most prominently in DNE,LOF and Impactful II *TP53* mutations. This effect was independent of presence or absence of *KRAS* mutations but was partially reduced in tumors with *TP53*-*EGFR* co-mutations. Hypoxia has been shown to induce EMT through *HIF-1α* as well as other mediators^92,93^ and to positively select for p53-deficient cells that have lost apoptotic potential in mice^94^. Thus, a hypoxic TME may provide a survival advantage for highly plastic *TP53*^mut^ LUAD malignant cells, which have lost alveolar identity and experience increased proliferative capacity and metastatic potential. This model is supported by our analysis of data from genetically engineered K and KP mouse models, and from A549 LUAD cell line (*TP53*^WT^) overexpressing different *TP53* mutational variants. Furthermore, *de novo* discovery of spatial multicellular communities revealed a highly hypoxic, EMT-promoting spatial niche enriched for *SPP1*^+^ TAMs, myofibroblasts, and collagen producing CAFs. *SPP1* expression was highly upregulated in *TP53*^mut^ monocytes and TAMs and tightly linked with regulation of hypoxia and EMT. Increased TGF-β signaling likely also helps shape an EMT-supporting niche in *TP53*^mut^ tumor samples, where *TGFB2*-*TGFBR2* interactions involving malignant cells were significantly enriched in spatial data. Previous studies linking mutant p53 to metastatic potential have focused on malignant cell-intrinsic processes^11,12,95^, as the effects of mutant p53 on tumor-stromal and tumor-immune crosstalk have been described only recently in model systems^96^. Our study is the first to profile the cellular and spatial context of *TP53* mutated LUAD tumors to provide insight into how *TP53* mutations could remodel the TME to promote tumor survival and metastasis in patients, leading to poorer outcomes.

Interestingly, one aspect of TME remodeling in *TP53*^mut^ LUAD is a higher tumor infiltration by B cells and *CD8*^+^ T cells. Increased lymphocyte recruitment may be caused in part by enriched *CXCL11*-*CXCR3* ligand-receptor interactions between myeloid and B/T cells in *TP53*^mut^ tumors, most notably exhausted T cells. We observed some overlap of the myeloid and T cell phenotypes in *TP53*^mut^ LUAD, the anti-tumor immunity hub we reported previously in colorectal cancer^97^, and the stem immunity hub in NSCLC^98^, but additionally report here B/Plasma cell tumor infiltration, T cell exhaustion (supported by an independent scRNA-seq validation compendium), and the enrichment of *SPP1*-expressing TAMs. Our findings of a potentially more immunogenic TME in *TP53*^mut^ LUAD is consistent with previous studies reporting higher levels of anti-tumor immune signatures^99^ in *TP53*^mut^ *vs*. *TP53*^WT^ LUAD using bulk RNA-seq data from TCGA and longer survival in advanced *TP53*^mut^ *vs*. *TP53*^WT^ NSCLC patients receiving immune checkpoint therapy^100^. Previous studies have reported that p53 mutations increase expression of PD-L1 *via* decreased inhibition by miR-34^101^ and that LUAD tumors with *KRAS*-*TP53* co-mutations have higher expression of PD-L1 and improved response to anti-PD1 treatment compared to *KRAS*^mut^*TP53*^WT^ tumors^19,21,23,102^. The link between mutation status and immunotherapy response has also been shown in the context of other mutations, where *KRAS*^mut^*STK11*^mut^*KEAP1*^mut^ LUAD tumors demonstrate reduced response compared to *KRAS*^mut^*STK11*^WT^*KEAP1*^WT^ tumors^103^. In agreement, our computational screen for immune checkpoint receptor-ligand pairs showed enrichment in *PDCD1*-*CD274*, *CTLA4*-*CD86*, and *TIGIT*-*PVR* interactions in *TP53*^mut^ *vs*. *TP53*^WT^ LUAD tumors; *CTLA4*-*CD86* and *TIGIT*-*PVR* interactions have not yet been studied in the context of *TP53* mutations to the best of our knowledge. Increased expression of *TIGIT* in T cells and *PVR* in malignant cell both contribute to the spatially-resolved enrichment of *TIGIT*-*PVR* co-expression, which we observed in all 8 *TP53*^mut^ LUAD tumors in our cohort, as well as in bulk RNA-seq data from TCGA and bulk proteomic data from CPTAC. Recent trials demonstrate benefit to combining anti-PD1 ICI with chemotherapy as a neo-adjuvant in resectable LUAD^104^. Moreover, trials targeting TIGIT alone or in combination with PD1/PD-L1 in lung cancer have yet to demonstrate substantial survival benefit, highlighting the importance of identifying subsets of patient tumors that could benefit from therapeutic intervention (*e.g. TP53*^mut^, PVR-expressing tumors)^105^. Future studies will be needed to assess the predictive potential of *TP53* mutational status for response to these and other therapies.

More generally, the methodology introduced in this study can be used to study the effect of *TP53* or other mutations on the TME in lung cancer and other tumor types and to infer multicellular pathways and communities to target for tumor suppressors in other contexts, where more effective therapies are needed.

## METHODS

### Human subjects

Fresh solid primary tumor tissue was collected using an IRB-approved protocol (IRB protocol 98-063) at DFCI/BWH and MGH. Patients were all confirmed to be treatment-naïve and consented to the study prior to collection. Immediately after resection, the tissue was reviewed by the clinical pathology team and high-quality portions (determined based on tumor content, necrosis, calcification, fat, and hemorrhage) were allocated for WES, scRNA-seq, spatial transcriptomics, and mIF. The protocol was designed to reduce the time between surgical resection, anatomic pathology review, placement in media, and processing at the Broad Institute. Blood was drawn from the same patients and cryo-preserved at -80 °C for subsequent processing. Patient demographic information, including age, sex, and smoking history, are included in **Table S1**.

### Whole-exome sequencing sample processing

DNA was extracted from fresh frozen lung cancer tissue embedded into OCT (TissueTek, Sakura) and from PBMCs isolated from preserved patient-matched blood (AllPrep DNA/RNA extraction kit, Qiagen). Library construction was performed as previously described^106^. In brief, DNA input for shearing was diluted to a final concentration of 20–250 ng in 50 µl of solution. Adapter ligation was performed using palindromic forked adapters (Integrated DNA Technologies), containing unique dual-indexed molecular barcode sequences to improve downstream pooling. End repair/A-tailing, adapter ligation, and library enrichment PCR were carried out using a 96-reaction kit format (Kapa HyperPrep). During solid-phase reversible immobilization (SPRI), a final elution volume of 30 µl was produced to maximize library concentration, and vortexing was performed to maximize the effluent. Constructed libraries were first pooled and hybridization and capture were performed (Illumina’s Nextera Exome Kit) using the recommended protocol from the manufacturer, using a skirted PCR plate to facilitate automation (Agilent Bravo liquid handling system). Library pools then underwent qPCR quantification and libraries were adjusted to a concentration of 2nM. DNA libraries were cluster-amplified using exclusion amplification chemistry in patterned flow cells (Illumina) according to the manufacturer’s recommended protocol. Sequencing of flow cells was performed using sequencing-by-synthesis chemistry and analyzed using RTA v.2.7.3 or later. Library pools were sequenced on 76 cycle runs using two 8-cycle index reads across the appropriate number of lanes to attain coverage for all libraries.

### Tissue processing, CD45 sorting, and scRNA-sequencing

Tumor tissue resections were transferred in RPMI on ice from the operating room and washed in cold PBS in the laboratory and transferred to 5ml Eppendorf tubes containing 3ml dissociation mixture (NSCLC PDEC protocol^107^). Samples were minced in the Eppendorf tube using scissors into pieces smaller than ∼0.4 mm and incubated at 37 °C, rotating at ∼14 rpm. for 10 min. Each sample was pipetted 20 times with a 1ml pipette tip at room temperature and placed back into incubation for 10 min. The sample was pipetted again 20 times using a 1 ml pipette tip, transferred to a 1.7 ml Eppendorf tube and centrifuged at 300–580*g* for 4–7 min at 4 °C. The pellet was resuspended in 200–500 µl of ACK (ammonium-chloride-potassium) RBC lysis buffer (Thermo Fisher Scientific, A1049201) and incubated for 1 min on ice. Twice the volume of cold PBS was added to stop the reaction, and cells were pelleted by a short centrifugation for 8 s at 4 °C, using the short spin setting and ramping up to 11,000*g*. If RBCs remained, the RBC lysis step was performed up to two additional times. To assess cell count and viability, 5 µl of Trypan blue (Thermo Fisher Scientific, cat. no. T10282) was mixed with 5 µl of the sample and loaded onto an INCYTO C-Chip Disposable Hemocytometer, Neubauer Improved (VWR, cat. no. 82030-468). Depletion of CD45^+^ cells in NSCLC samples was performed using CD45 MicroBeads (Miltenyi Biotec, cat. no. 130-045-801) according to the manufacturer’s protocol. The single-cell suspension was centrifuged at 500*g* for 4 min at 4 °C and the pellet was resuspended in 80 µl of MACS buffer (PBS supplemented with 0.5% BSA, and 2 mM EDTA) per 10^6^ cells. MACS CD45 microbeads were added to the cell suspension (20 µl per 10 million cells) and the cells were incubated on ice for 15 min. During incubation, a MS column was attached to a MidiMACS separator and rinsed with 3 ml MACS buffer. After incubation, the cells-bead conjugate was washed with 900 µl MACS buffer per 10 million cells. The cells were then centrifuged at 500*g* for 4 min at 4 °C and the pellet was resuspended in 500 µl MACS buffer. The cell suspension was transferred to the column and the effluent was collected (CD45^−^ fraction). The column was washed three times with 3 ml MACS buffer. The CD45^−^ fraction was centrifuged at 500*g* for 4 min at 4 °C. The pellet was resuspended in 50 µl of 0.4% BSA (Ambion, cat. no. AM2616) in PBS. Cells were re-counted and adjusted to a final range of 200–2,000 cells per µl.

### NSCLC tissue dissociation protocol

The digestion mix contained 2692 µl HBSS (Thermo Fisher Scientific, cat. no. 14170112), 187.5 µl of 20 mg ml^−1^ pronase (Sigma Aldrich, cat. no. 10165921001) diluted to a final concentration of 1,250 µg ml^−1^, 27.6 µl of 1 mg ml^−1^ elastase (Thermo Fisher Scientific, cat. no. NC9301601) diluted to a final concentration of 9.2 µg ml^−1^, 30 µl of 10 mg ml^−1^ DNase I (Sigma Aldrich, cat. no. 11284932001) diluted to a final concentration of 100 µg ml^−1^, 30 µl of 10 mg ml^−1^ Dispase (Sigma Aldrich, cat. no. 4942078001) diluted to a final concentration of 100 µg ml^−1^, 30 µl of 150 mg ml^−1^ collagenase A (Sigma Aldrich, cat. no. 10103578001) diluted to a final concentration of 1,500 µg ml^−1^ and 3 µl of 100 mg ml^−1^ collagenase IV (Thermo Fisher Scientific, cat. no. NC9836075) diluted to a final concentration of 100 µg ml^−1^.

### scRNA-seq library preparation and sequencing

A total of 8,000 cells were loaded onto each channel of the Chromium Controller (10X Genomics) to generate single-cell gel bead-in-emulsions (GEMs). Libraries were constructed using the Chromium Single Cell 3ʹ Library & Gel Bead Kit v.2 or v.3 (PN-120237, 10X Genomics): barcoded reverse transcription of RNA, cDNA amplification, fragmentation and adapter and sample index attachment were all performed according to the manufacturer’s recommended protocol. Barcoded libraries from four 10x channels were pooled together and sequenced on either one lane of an Illumina HiSeq X or one flow cell of a NextSeq, with paired end reads accordingly: read 1, 26 nt; read 2, 55 nt; index 1, 8 nt; index 2, 0 nt.

### Tissue processing for Spatial Transcriptomics

Fresh frozen lung cancer tissue samples were embedded into OCT (TissueTek, Sakura) and shipped to the Broad Institute for cryosectioning and generation of hematoxylin and eosin-stained (H&E) tissue sections. H&E-stained tissues were subject to pathology review to assess tissue quality, structural preservation, cellular viability, tumor content, inflammation, fibrosis, and necrosis. Samples were excluded based on small tissue size, low cellularity, or extensive fibrosis. Regions of interest (ROIs) were then selected based on the criteria mentioned and marked on H&E images for subsequent tissue sectioning and mounting on Visium slides. Visium sectioning and processing were performed on the eight samples that passed QC. Tissues were cryosectioned at 10 um thickness at -22°C and placed in the capture areas of Visium Tissue Optimization slides (3000394, 10X Genomics) and Visium Spatial Gene Expression slides (2000233, 10X Genomics). The tissue sections were adhered by warming the backside of the slides and placed at -80°C for 1-3 days.

### Visium spatial gene expression library generation

The tissue optimization sample slide and spatial gene expression slide were processed according to the manufacturer’s protocols (CG000238_VisiumSpatialTissueOptimizationUserGuide_Rev_A.pdf and CG000239_VisiumSpatialGeneExpression_UserGuide_Rev_C.pdf). In short, following tissue methanol fixation and H&E staining, brightfield morphology images were obtained with a Zeiss Axio microscope using the Metafer slide-scanning platform (Metasystems) at 10x resolution.

The images were joined together with the VSlide software (Metasystems) and exported as tiff files. Optimal tissue permeabilization times were tested with eight different time points (0, 3, 6, 12, 18, 24, 30, 36 minutes) on the tissue optimization sample slide for one of the samples. 12 minutes of permeabilization was set as the optimal time point and used for the spatial gene expression slide for all samples. The released RNAs from the tissue were reversely transcribed into complimentary DNA by priming to the spatial barcoded primers on the glass in the presence of template switching oligo. After second strand synthesis, a denaturation step released the cDNAs, which were then amplified with 14-15 cycles of PCR amplification. Finally, indexed sequencing-ready, spatial gene expression libraries were constructed according to the manufacturer’s protocol. The libraries were pooled together to generate >50,000 reads per spatial spot and sequenced on Illumina NovaSeq sequencers with S1 and SP kits and the settings: read 1, 28 cycles; read 2, 90 cycles; index 1, 10 cycles; index 2, 10 cycles.

### Multiplex immunofluorescence

Staining was performed on a BOND RX fully automated stainer (Leica Biosystems). FFPE tissue sections of 5-μm thick were baked for 3 hours at 60°C before being loaded into the BOND RX. Slides were deparaffinized (BOND DeWax Solution, Leica Biosystems, Cat. AR9590) and rehydrated with multiple rounds of graded ethanol to deionized water. Antigen retrieval was performed in BOND Epitope Retrieval Solution 1 (pH 6) or 2 (pH 9) (ER1, ER2, Leica Biosystems, Cat. AR9961, AR9640) at 95°C. Deparaffinization, rehydration and antigen retrieval are carried out by BOND RX for ∼ 14 hours. Slides were then serially stained with primary antibodies with an incubation time of 30 minutes per antibody. Anti-mouse and anti-rabbit Opal Polymer Horseradish Peroxidase (Opal Polymer HRP Ms + Rb, Akoya Biosciences, Cat. ARH1001EA) were then applied as a secondary label for 10 minutes. Signal for antibody complexes was labeled and visualized by their corresponding Opal Fluorophore Reagents (Akoya) by incubating the slides for 10 minutes. Opal Fluorophore 780 was paired with a TSA-DIG amplification to ensure analyzable signal. Slides were incubated in Spectral DAPI solution (Akoya) for 10 minutes, air dried and mounted with Prolong Diamond Anti-fade mounting medium (Life Technologies, Cat. P36965) and stored in a light-proof box at 4 C° prior to imaging.

Imaging was performed using the PhenoImager multispectral imaging platform (Akoya Biosciences, Marlborough, MA). Each slide was scanned at 20x resolution as a whole-slide image. These images were then accessed through Phenochart viewing software (Akoya Biosciences) where 4-8 20x regions of interest (ROIs) were selected. After ROI selection was approved by a pathologist (S.J.R.), the images were spectrally unmixed and analyzed using Inform 2.6 (Akoya Biociences). Each analyzable ROI was segmented and quantified for expression of each marker utilizing the Inform analysis tools.

### WES data processing

The Picard pipeline (http://picard.sourceforge.net/) was used to align the tumor and normal whole exome sequences to the hg19 reference human genome build and generate BAM files. Somatic short variant discovery was performed using the Mutect 2 algorithm (https://software.broadinstitute.org/gatk/documentation/tooldocs/current/org_broadinstitute_gatk_tools_walkers_cancer_m2_MuTect2) from the GAT4K pipeline run on the Terra cloud-based platform. Somatic mutations were annotated using Funcotator (https://gatk.broadinstitute.org/hc/en-us/articles/360035889931-Funcotator-Information-and-Tutorial). Copy number ratios for each exon was calculated by comparing mean exon coverage with expected coverage based on a panel of normal samples and were then segmented for downstream analysis. Maf and vcf files were generated as outputs, which were then subsequently processed with maftools^108^ and cnvkit^109^ packages.

### Large-scale CNA inference and correlation

The inferCNV package^110^ (https://github.com/broadinstitute/inferCNV) was applied to infer large-scale CNVs from scRNA-seq data. Normal lung epithelial cells from a separate dataset were used as a reference. Raw UMI counts, a gene ordering file, and an annotation file specifying normal and tumor cells and patient sample of origin were used as input. The function infercnv::run() was performed with a cutoff of 0.1, cluster_by_groups = T, and HMM = T. CNV values were predicted using a 6-state Hidden Markov Model (HMM) and were normalized to the 3^rd^ quantile of all values per sample. Cells with > 0.7 normalized CNV values were assigned as malignant cells. CNVs called from WES data were used as a positive control to assess the accuracy of the inferCNV analysis using scRNA-seq data. Spearman correlation was run between inferred CNVs from scRNA and from WES data across genes and plotted as a heatmap.

### scRNA-seq preprocessing

CellRanger v3.1 was used to align 10X Chromium reads to the GRCh38 human genome reference and generate barcode, gene, and count matrices. Preprocessed matrices were then loaded into Seurat v4^111^ in R v4.1.0 using the Read10X() function. Cells with less than 1000 unique molecular identifiers (UMIs), less than 400 detected genes (1000 genes for malignant cells), or greater than 25% mitochondrial genes were excluded from further analyses.

### scRNA-seq integration, clustering, annotation, and doublet removal

The UMI counts matrices were batch-corrected by patient sample, using the Seurat v4 SCTransform() (using 2000 variable features and regressing out percentage of mitochondrial genes), FindIntegrationAnchors(), and IntegrateData(), using the largest sample as the reference dataset. PCA was then performed in the variable gene space and 15 principal components were used for Louvain clustering and UMAP dimensionality reduction. Cell type markers for broad cell classes were identified using the FindAllMarkers() function, using a maximum of 1,000 cells per cluster. Cells were annotated based on expression of canonical markers that were established by previous literature. Individual cell classes were then subset and subjected to another series of integration, clustering, and annotation based on expression of subset-specific markers. Doublets were removed based on expression of markers specific to more than one broad cell classes.

### scRNA-seq signature scoring

To identify broad cell classes signatures, we used the FindAllMarkers() function in Seurat v4 and generated gene sets using the most differentially expressed genes for each class, based on log_2_ fold change and Bonferroni-corrected *p*-values compared to the other clusters. We identified malignant program signatures by performing anchor-based integration across patient samples malignant cells followed by Louvain clustering at resolution 1.2 and using the FindAllMarkers() function to derive the top 10 most differentially expressed genes from each cluster, based on log_2_ fold change and Bonferroni-corrected *p*-values. Signatures for normal lung epithelial cells were derived from published healthy lung scRNA-seq data using the same method. The lists of broad cell classes and cell subset marker genes, as well as final gene set signatures are provided in **Table S2**. To define hallmark EMT and hypoxia scores, we used gene sets termed “HALLMARK_EPITHELIAL_MESENCHYMAL_TRANSITION” and “HALLMARK_HYPOXIA” downloaded from MSigDB^112^. Healthy lung epithelial cell signatures were derived using published scRNA-seq data from the human lung cell atlas^113^, as well as *a priori* knowledge: AT2 (*SFTPC*, *SFTPA2*, *SFTPA1*, *SFTPD*, *SFTPB*), AT1 (*AGER*, *EMP2*, *CAV1*, *HOPX*), basal (*KRT5*, *KRT17*, *KRT15*, *S100A2*), neuroendocrine (*CALCA*, *CHGA*, *GRP*, *SCG5*, *ASCL1*), goblet (*MUC5AC*, *MUC5B*, *S100P*), secretory (*BPIFB1*, *PIGR*, *SCGB3A1*, *SCGB1A1*), ciliated (*TMEM190*, *CAPS*, *FOXJ1*, *TPPP3*, *FAM183A*), and ionocyte (*ASCL3*, *ATP6V0B*, *FOXI1*, *CLCNKB*, *CFTR*). Signature scores were assigned per cell in scRNA-seq data using the AddModuleScore() function in Seurat v4, which compares the average expression of the genes in each gene set to the average expression of a control gene set of 100 randomly selected genes within the same expression bin.

### scRNA-seq differential expression analysis

The data was first split into different cell subsets, and pseudobulk profiles were generated by setting the identity of the objects to patient sample ID and running the AverageExpression() function in Seurat v4. A Mann-Whitney Wilcoxon test was used to test for Bonferroni-corrected significance in gene differential expression between groups. The average log_2_ fold change was plotted using the ComplexHeatmap package in R, and asterisks were added for genes that were significantly differentially expressed.

### Gene Set Enrichment Analysis

Using the top 100 DEGs between conditions per cell subset or the top 100 specific markers per subset, we performed gene-set enrichment analysis using the R package clusterProfiler^114^ (v4.0), focusing on GO terms, KEGG, Reactome, and Hallmark Cancer gene sets. Gene ratios and adjusted *p*-values were visualized as dot plots, after applying BH correction and setting *p*-value and *q*-value cutoffs of 0.05.

### Similarity to normal lung and K/KP mouse cells

To identify transcriptomic similarities between cell subsets in NSCLC and the normal lung, the Seurat v4 function TransferData() was used. The broad cell compartment object in the NSCLC scRNA-seq dataset was assigned to the query parameter and the corresponding compartment in a published normal lung scRNA-seq dataset^56^ was assigned to the reference parameter, and the remaining parameters were set to defaults. Each cell in the NSCLC object was assigned a predicted identity based on the normal lung subset it was most similar to transcriptionally. A confusion matrix was created, displaying the number of cells that overlapped between predicted identities from the normal lung and annotations assigned *de novo*. A similar approach was used to classify K/KP mice HPCS cells to their closest annotation in human tumor cells.

### Signaling Entropy calculation

Entropy was inferred for each epithelial cell using the R package SCENT^64^, using the default protein-protein interaction (PPI) network (“net17Jan2016.m”). Raw UMI counts were used as the scRNA-seq input. Entropy was calculated separately for malignant and non-malignant epithelial cells. Quality control was performed using the scater package in R, and low-quality cells based on library size, number of expressed genes, and mitochondrial gene proportion were removed. Differential potency estimation was performed using the DoIntegPPI() function, which generated entropy values per cell. The same method was used to calculate entropy values for bulk RNA-seq data from TCGA and bulk proteomic data from CPTAC. Entropy values were compared using the ggviolin() function using the ggpubr package in R.

### Cell-cell interaction analysis

CellPhoneDB^115^ (v2.1.7) was used to predict significant ligand-receptor interactions using normalized raw counts and annotation of broad cell classes as input. The method was run with default parameters separately for each sample ID to account for batch effects between samples. Interactions with a *p*-value below 0.05 were classified as significant. The significant means output was converted into a binary matrix based on presence or absence of a significant interaction. Normalized enrichment difference (*ED*) of a specific interaction was calculated as follows:

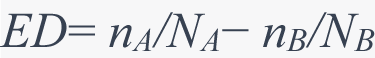

where *nA* and *nB* represent the number of tumor samples with the interaction in groups A and B, respectively, while *NA* and *NB* represent the total number of tumor samples in these groups.

Significance of enrichment for an interaction was calculated using a Fisher’s exact test. Negative log_10_ *p*-values and normalized enrichment differences for each ligand-receptor interaction and cell pair combination were plotted using the ggplot2 package in R.

### scRNA-seq regulon analysis

A scRNA-seq transcription factor (TF) gene regulatory network was constructed using pySCENIC^73^ (v0.11.2). Raw UMI counts were used as input and regulons were predicted using gene module co-expression using the GRNBoost2 package. The number of regulons was pruned using the feather ranking databases (hg19-500bp-upstream-7species.mc9nr, hg19-tss-centered-10kb-10species.mc9nr, hg19-tss-centered-5kb-10species.mc9nr). Regulon activity, driving TFs, and weights for individual target genes were predicted with the cisTarget() function in pySCENIC. Regulon activity enrichment scores were predicted for each cellular subset using AUCell and regulon specificity scores were used to identify regulatory networks specific to each cell subset. Z-score-transformed regulon x subset matrices were visualized using the ComplexHeatmap package in R.

### Additional scRNA-seq datasets

Processed scRNA-seq data from an additional 5 studies (both published and unpublished) with available *TP53* mutational status were assembled into a validation compendium, consisting of 15 *TP53*^WT^ and 7 *TP53*^mut^ tumor samples (ArrayExpress accessions E-MTAB-6149 and E-MTAB-6653^26^; https://humantumoratlas.org/ (unpublished); GEO accession GSE123904^32^; BIG accession HRA000154^55^; https://doi.org/10.24433/CO.0121060.v133). *In*-*vivo* scRNA-seq data for *KRAS*^mut^*TP53*^WT^ and *KRAS*^mut^*TP53*^mut^ mice (Smart-Seq2) was obtained from GEO accession GSE152607^63^. *In*-*vitro* scRNA-seq data from lung cancer A549 cell lines with different induced *TP53* variants was obtained from GEO accession GSE161824^61^.

### Signature scoring of bulk expression profiles

Bulk RNA-seq and somatic mutation data from TCGA LUAD and somatic mutations were downloaded from cBioPortal^116^ using the cgdsr package in R. Curated gene sets with highly specific markers for individual cell types, subsets, or malignant programs were generated from the scRNA-seq data as described above. The signature scores for these gene sets were defined as the mean log_10_ expression of all the genes in a given gene set and were used to estimate the relative frequency of cell types, cell subsets in bulk RNA-seq data from TCGA, or activity of malignant cell programs. The same approach was used for calculating expression signatures in bulk proteomic data from CPTAC^117^, which was downloaded from the CPTAC Data Portal https://cptac-data-portal.georgetown.edu/cptacPublic/. We confirm the reproducibility of all our bulk deconvolution results using BayesPrism, a deconvolution method based on a Bayesian framework that incorporates single-cell expression profiles^118^.

### Survival analysis

To predict the roles of individual genes and gene signatures on patient survival, TCGA LUAD tumor samples were classified into high (top 25^th^ percentile), medium (25^th^ to 75^th^ percentile), and low (bottom 25^th^ percentile); or high (top 50^th^ percentile) and low (bottom 50^th^ percentile) expression groups. Kaplan-Meier (KM) curves were generated using the survival package in R, and *p*-values were calculated using a log-rank test or bivariate Cox proportional-hazards regression analysis to evaluate the significance of different model parameters (*e*.*g*., expression level of genes or malignant signatures, *TP53* mutation status) in relation to overall survival (defined as time to death from any cause). The reported *p-*values from the Cox proportional-hazards model reflect the significance of one specific covariate while adjusting for *TP53* mutation status.

### Effects of co-mutations and *TP53* variant classes

TCGA tumor samples were stratified into six mutually exclusive categories with sufficiently large sample sizes for comparative analysis based on combinations of mutations in *KRAS*, *EGFR*, and *TP53* (*KRAS*^mut^, *EGFR*^mut^, *TP53*^mut^, *KRAS*^mut^*TP53*^mut^, *EGFR*^mut^*TP53*^mut^, and *KRAS*^WT^*EGFR*^WT^ *TP53*^WT^). TCGA and CPTAC tumor samples were furthermore grouped into different mutually exclusive missense variant classes based on either dominant-negative/loss of function (DNE,LOF) status of mutational variants, or impact categories as described by prior studies^60,61^. Mann-Whitney-Wilcoxon test *p*-values were computed for each group relative to the relevant wildtype or control group.

### Multiple linear regression and ANOVA analysis of interaction effects

Expression level of individual genes or gene signatures was predicted based on the mutation status of four different genes (*EGFR*, *KRAS*, *STK11*, and *TP53*), and tumor mutational burden (TMB) using the following model:

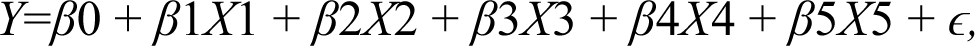

where *Y* represents the expression level, *β*0 the y-intercept, *β*1, *β*2, *β*3, *β*4, and *β*5 the regression coefficients associated with each variable, *X*1, *X*2, *X*3, and *X*4 the mutation status of different genes (*EGFR*, *KRAS*, *STK11*, and *TP53*), *X*5 the TMB, and *ɛ* the error term. For each mutation variable *Xi*, a *p*-value was computed to test the null hypothesis that the regression coefficient *βi* is equal to zero (indicating no effect). A significant *p*-value (≤ 0.05) suggests that the mutation status of the respective gene is a significant predictor of *Y* (*i.e.,* there exists a significant relationship between the mutation status and the expression level). We further conducted the same analysis using a model containing an expanded set of covariates (mutation status of *EGFR*, *KRAS-G12C*, *KRAS-G12D*, *KRAS-G12V*, *STK11*, *KEAP1*, *RBM10*, *PTPRD*, and *TP53* as well as TMB). An analysis of variance (ANOVA) was conducted on the output of the regression model’s output, evaluating the interactions between *KRAS* and *TP53*, and between *EGFR* and *TP53*. A significant interaction implies that the combined effect of two individual interactions is not merely the sum of their individual effects. The output from the ANOVA provides *p*-values for each interaction, where a significant *p*-value suggests that the effect of one mutation is altered by the presence or absence of the other mutation (*i.e.,* not additive).

### Pathologist annotation of hematoxylin and eosin (H&E) stains from spatial transcriptomics samples

A board-certified pathologist, blinded to all sample identities and spatial transcriptomics measurements, performed manual annotation of 13 H&E stains using Loupe browser.

Annotations on the H&E were classified into multiple categories (*e.g*., cancer, lymphocytes, myeloid, fibroblast, lepidic adenocarcinoma, TLS, vascular endothelium) that were most representative of cells within barcoded spots. Spots in which there were cell types whose identities could not be readily visually resolved were not annotated. These categories were then mapped to three general cell classes (malignant, stromal, and immune) to minimize variability in pathology annotations across samples and allow for comparison with the computationally inferred cellular compositions.

### Processing of spatial transcriptomics data, integration with paired scRNA-seq data, and colocalization analysis

Tangram version 0.4.0, scanpy 1.8.1, and anndata 0.7.6 was used to map scRNA-seq data to spatial locations on 10X Visium spatial transcriptomics samples as previously described^41^. UMI counts for scRNA-seq and spatial transcriptomics data were converted into anndata objects and pre-processed using standard scanpy functions^119^. The top 100 genes that best characterized each broad cell classes were selected by sc.tl.rank_genes_groups(). This subset of genes was used afterwards as training genes for Tangram alignment. To more accurately map single-cells onto spatial transcriptomics spots, an estimate of cell density per spot was attained through watershed-based nuclei segmentation of the hematoxylin and eosin-stained serial tissue section. Images were first imported by squidpy 1.1.0^120^, and smoothened using squidpy.im.process with a sigma = 4, then segmented by squidpy.im.segment with method = “watershed”, using parameters channel = 0 to select the red color channel and thres = 120. A threshold of 120 was selected to best separate nuclei from background, and sq.im.calculate_features was used to count the number of segmented nuclei per spot after segmentation. Finally, mapping was performed by tg.map_cells_to_space using mode = “constrained” and density_prior = # of nuclei per spot / total # of nuclei to constrain alignment based on segmented nuclei density, returning probabilities of spatial location on a per-cell basis, which were employed in later analysis. To project individual gene expression onto spatial transcriptomics spots, tg.project_genes was used. Cell type and subset proportion were predicted per spot by summing up cell x spot probabilities and normalizing the total probability scores per spot to 1. Spatial colocalization was inferred using Pearson correlation between predicted cell type and subset compositions, and additional continuous attributes from the metadata across spots.

### NMF analysis for spatial transcriptomics data

NMF analysis was adapted from cell2location^121^ using cell subset proportions per spatial transcriptomics spot as inputs, enabling analysis of spatially colocalized cellular programs. NMF was trained five times for a range of k = 3 to 30 factors, and k = 14 was selected based on stability of training across the five restarts and elucidating discrete spatially-delimited compartments across spatial transcriptomics samples. Using scanpy, NMF weights were plotted onto spatial transcriptomics data to investigate spatial-dependent patterning of NMF factors.

NMF weights per subset were visualized using the heatmap function in cell2location.

### Multiplex immunofluorescence analysis

InForm-analyzed ROIs were processed using the Pythologist software^122^. Cell colocalization involved three spatial categories: 1) direct contact, 2) adjacency within 6 nearest neighbors, and 3) proximity within 50um of PD-L1^+^cytokeratin^+^ cells. Proportion of PD1^+^CD8^+^ cells out of all PD-L1^+^cytokeratin^+^ neighboring cells were computed across these spatially defined categories when at least 20 neighboring cells were present.

## ACKNOWLEDGMENTS

We thank Leslie Gaffney for extensive help in editing and graphical design of the figures in the manuscript, the Broad Genomics Platform, Broad Flow Cytometry Facility, Pathology and Surgery Departments at MGH and BWH; members of the Bueno, Rodig, Hata, Regev, Johnson, and Tsankov labs. The project is part of the Human Tumor Atlas Pilot Project (HTAPP) consortium and National Institutes of Health (NIH) HTAN (Human Tumor Atlas Network) consortium paper package. Work was supported by the Klarman Cell Observatory (to O.R.-R. and A. Regev) and also funded in part by federal funds from the National Cancer Institute (NCI), NIH Task Order HHSN261100039 under Contract HHSN261201500003, and U2CCA233195. A.M.T. was supported by Department of the Army Lung Cancer Research Program Career Development Award W81XWH2210079, American Cancer Society Research Scholar Grant RSG-23-1039063-01-MM, and ISMMS seed funding. W.Z. and B.Y.S. were supported by NIH grant 2T32GM007280-41 as part of the ISMMS Medical Scientist Training Program. The graphical summary was created with Biorender.com.

---

**Figure S1.**
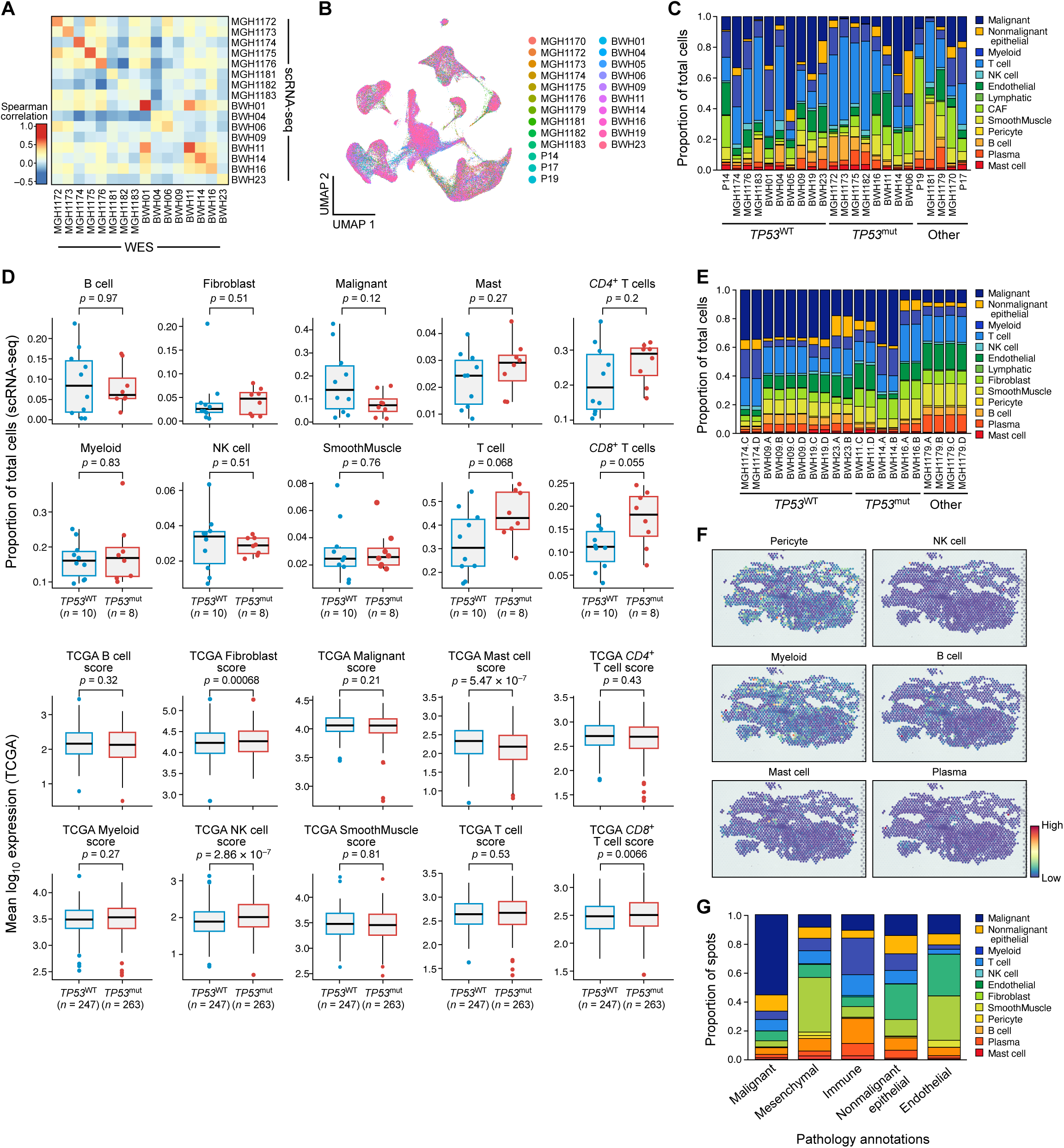
Integration and analysis of WES, scRNA-seq, and spatial transcriptomics data across tumors. **A.** Spearman correlation shows correspondence between the log_2_ CNA ratio predicted by scRNA-seq and WES data from the same patient. Correlation was computed across genes for each patient. Low correspondence in two tumor samples (MGH1183 and BWH09) likely resulted from small number of malignant cells (MGH1183) and intratumoral heterogeneity (BWH09). **B.** UMAP of 167,193 cells, colored by patient identity shows integration between tumor samples. **C.** Stacked bar plot displaying the composition of broad cell classes for each scRNA-seq patient sample. **D.** Top: Proportion of broad cell classes (not shown in Figure 1F) out of all cells found in the scRNA-seq data from *TP53*^WT^ (blue) and *TP53*^mut^ (red) LUAD tumors. Bottom: Average log_10_ expression of highly specific broad cell class markers derived from the scRNA-seq data for *TP53*^WT^ (blue) and *TP53*^mut^ (red) LUAD bulk RNA-seq tumor samples from TCGA. *P*-values were calculated using a Mann-Whitney-Wilcoxon test (scRNA-seq, top) or a two-tailed multiple regression t-test (TCGA, bottom). **E.** Stacked bar plot depicting mean cell composition across ST tumor samples, predicted using Tangram mapping. **F.** Spatial feature plots of broad cell classes not shown in Figure 1G, predicted by Tangram mapping. **G.** Mean proportion of Tangram-predicted cell classes (y-axis) across different manual spot annotations independently performed by a pathologist (NRM) without knowledge of sample and spot identities.

**Figure S2.**
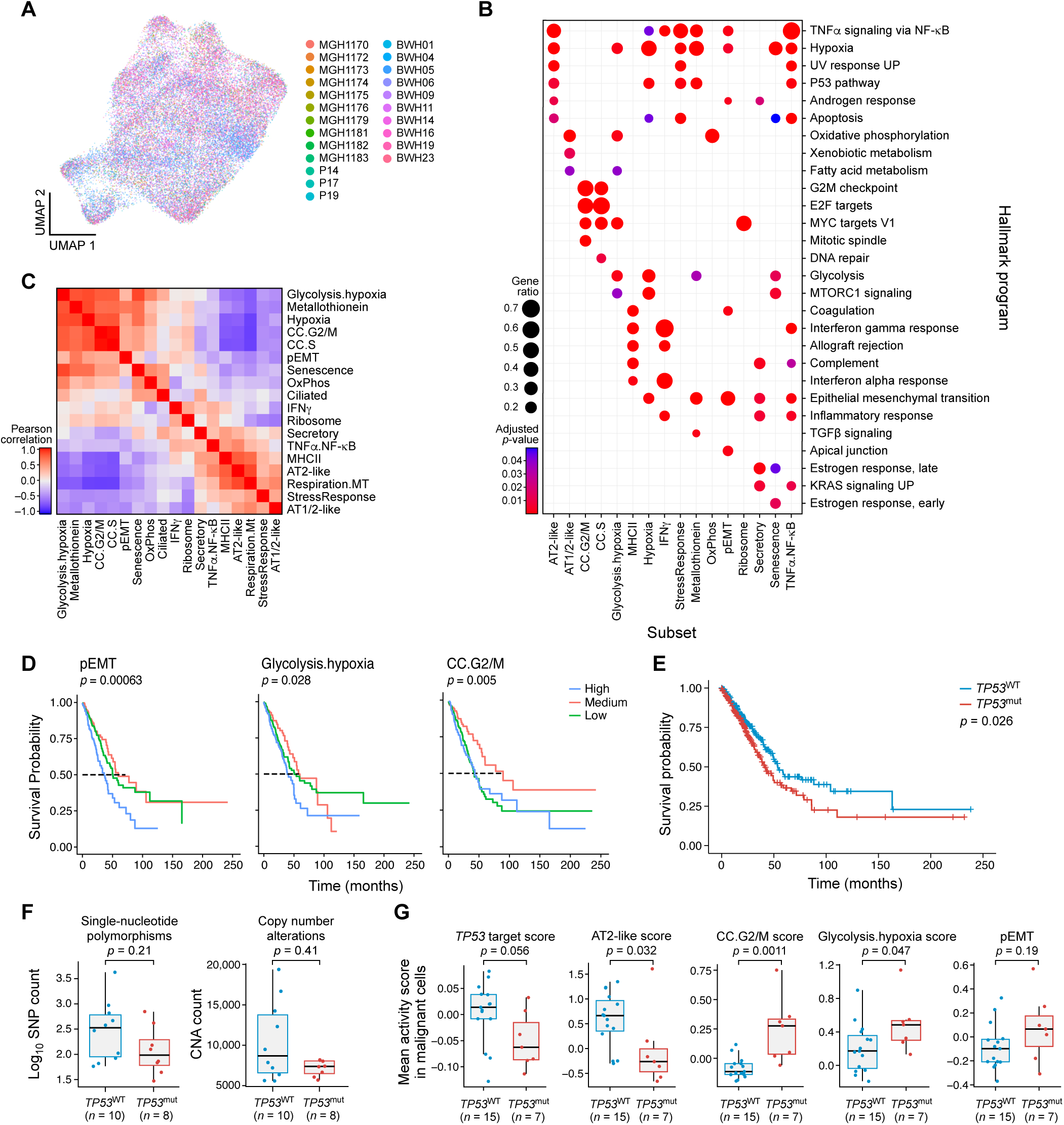
Characterization of the malignant compartment in NSCLC. **A.** UMAP of 33,377 malignant cells, integrated across tumors and colored by patient. **B.** Dot plot of gene set enrichment outputs (Hallmark) of 100 most differentially expressed markers for each malignant subcluster. **C.** Pearson correlation of average malignant program scores across tumors, analogous to Figure 2C but performed across patients instead of across cells. **D.** Kaplan Meier survival curves showing association of malignant program expression with survival in bulk TCGA LUAD. Tumors (n = 510) were stratified by program expression into high, medium, and low expression categories. Cox proportional-hazards model was used to calculate *p*-value corrected for *TP53* mutational status. **E.** Kaplan Meier survival curves showing association of *TP53* mutation status with survival in bulk TCGA LUAD. *P*-values were calculated using a log-rank test. **F.** Comparison of log_10_ SNP and CNA abundance between *TP53*^mut^ *vs. TP53*^WT^ tumors shows no significant difference in our cohort. **G.** Comparison of malignant program mean expression scores between *TP53*^mut^ and *TP53*^WT^ tumors in scRNA-seq validation cohort. *P*-values were calculated using a Mann-Whitney-Wilcoxon test.

**Figure S3.**
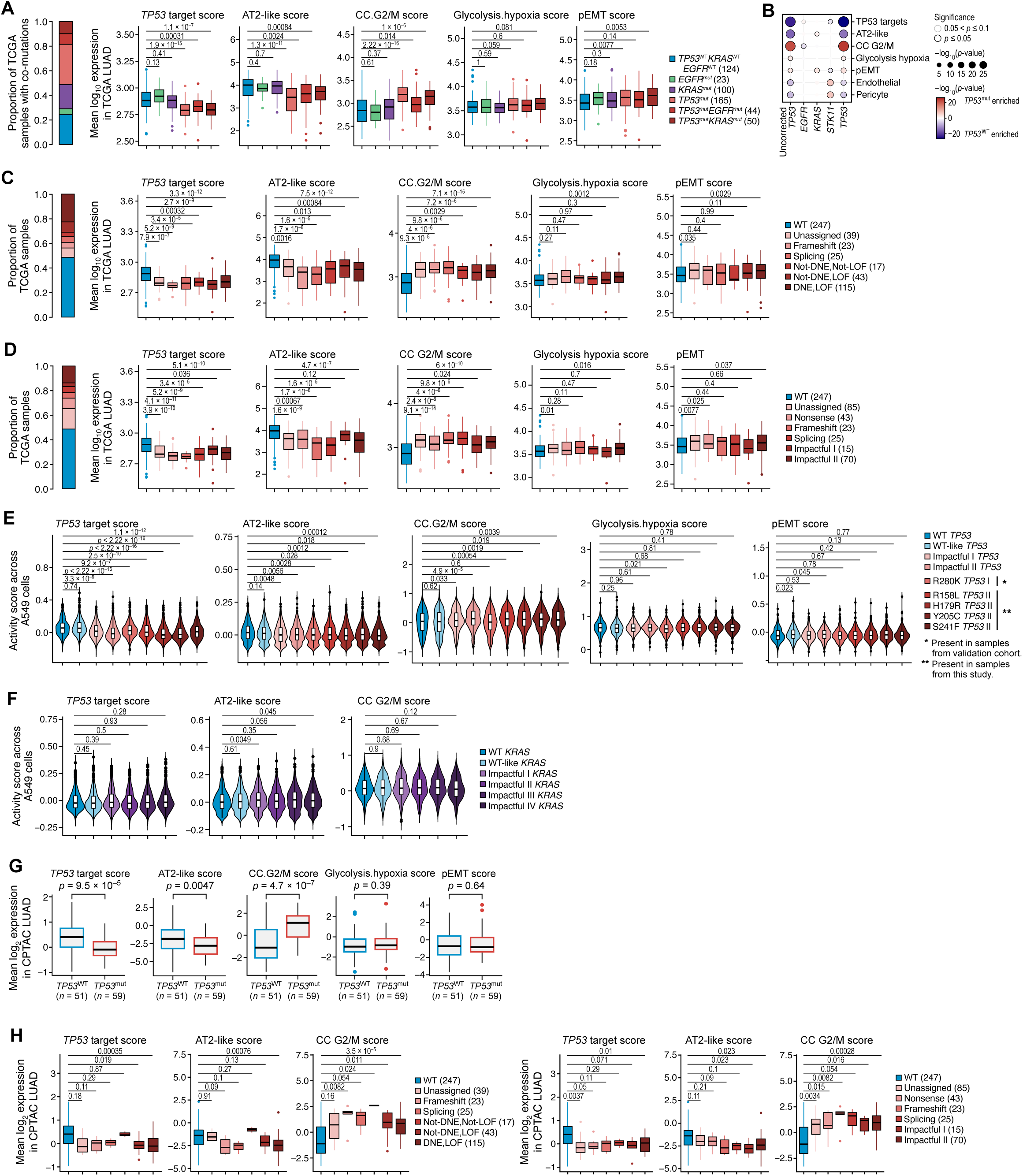
Consistent changes in malignant program expression and cellular entropy in *TP53*^mut^ LUAD across co-mutations, variant classes, cell lines, mouse models, bulk transcriptomic, and proteomic data. **A.** Left: Stacked bar plot of proportion of tumor samples with different co-mutations in TCGA. Right: Comparison of malignant program mean expression between *TP53*^WT^ and different single- or co-mutations in *EGFR*, *KRAS*, and *TP53* in bulk RNA-seq data from TCGA tumor samples. **B.** Dot plot showing association of common LUAD mutations (columns) with mean log_10_ expression of malignant programs, endothelial, and pericyte scores (rows) in TCGA LUAD. Color and size of dots correspond to -log10 *p*-value assessed by Mann-Whitney-Wilcoxon (uncorrected *TP53*, left) or a two-tailed multiple regression t-test (*EGFR*, *KRAS, STK11*, *TP53*). Black outline indicates *p*-value ≤ 0.05, and gray outline indicates *p*-value of ≤ 0.1. **C.** Left: Stacked bar plot of proportion of tumor samples with different categories of dominant-negative/loss of function *TP53*^mut^ effect in TCGA. Right: Comparison of malignant program mean expression between *TP53*^WT^ and different functional *TP53*^mut^ categories in TCGA tumor samples. *P*-values were calculated using a Mann-Whitney-Wilcoxon test. **D.** Left: Stacked bar plot of proportion of tumor samples with different impact categories in TCGA. Right: Comparison of malignant program mean log_10_ expression between *TP53*^WT^ and different functional *TP53*^mut^ impact categories in TCGA tumor samples. *P*-values were calculated using a Mann-Whitney-Wilcoxon test. **E.** Comparison of malignant program mean expression between *TP53*^WT^ and different *TP53*^mut^ impact categories^59^ in A549 cells. Rightmost four violin plots in each panel show the effect of mutations also found in tumor samples from our cohort. **F.** Comparison of malignant program mean expression scores between *KRAS*^WT^ and different *KRAS*^mut^ impact categories in A549 cells. **G.** Comparison of malignant program mean expression between *TP53*^mut^ and *TP53*^WT^ tumors using bulk proteomic data from CPTAC. *P*-values were calculated using a Mann-Whitney-Wilcoxon test. **H.** Comparison of malignant program mean log_2_ expression between *TP53*^WT^ and different dominant-negative/loss of function (left) and impact categories^59^ (right) of *TP53*^mut^ tumor samples in bulk proteomic data from CPTAC. *P*-values were calculated using a Mann-Whitney-Wilcoxon test.

**Figure S4.**
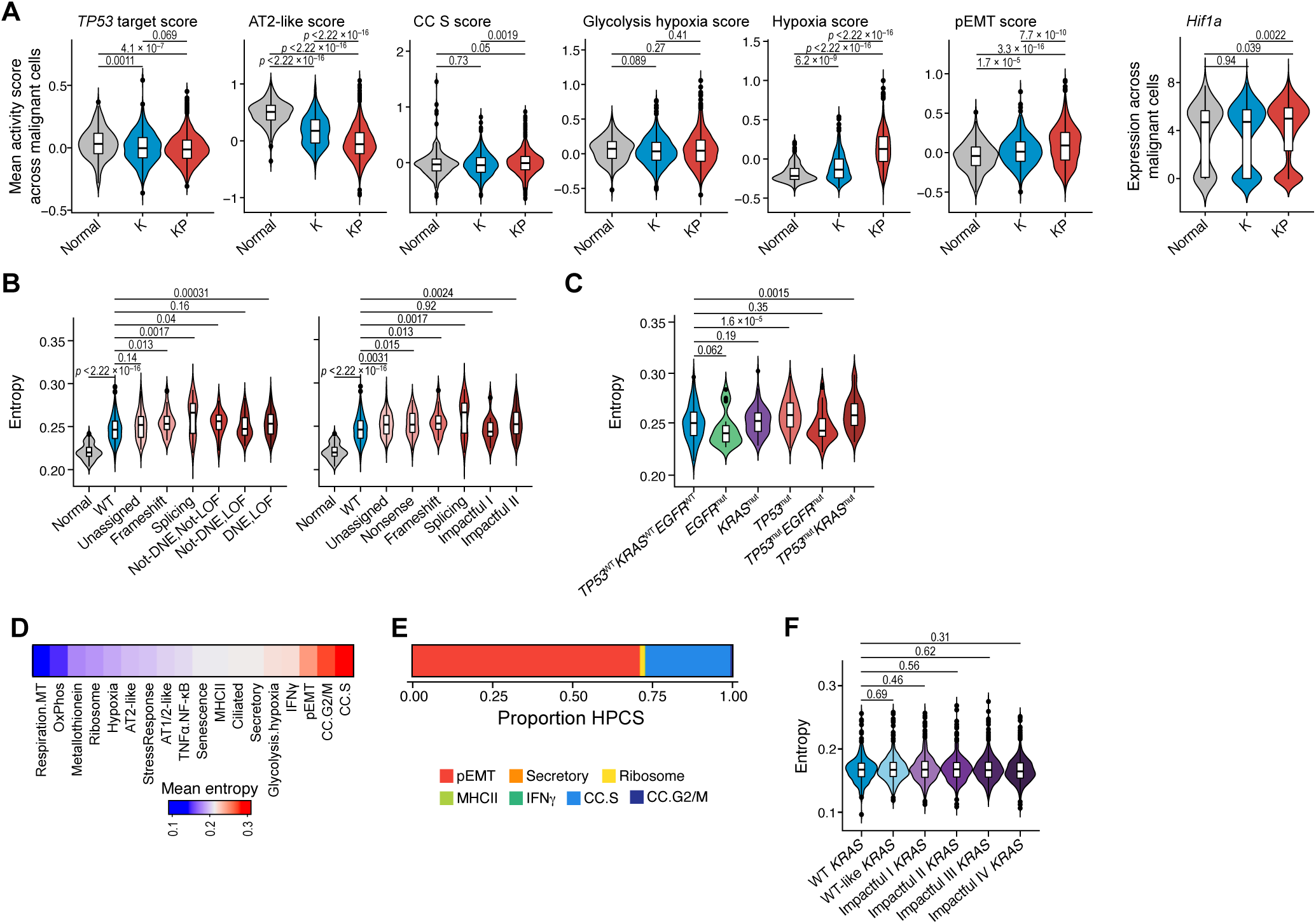
Effect of different co-mutations and *TP53* mutation variants on malignant program activity and signaling entropy. **A.** Comparison of malignant program scores and *Hif1a* expression between K (blue) *vs.* KP (red) LUAD mouse model scRNA-seq data malignant cells. *P*-values were calculated using a Mann-Whitney-Wilcoxon test. **B.** Distribution of entropy scores across adjacent normal (grey), *TP53*^WT^ (blue), and different *TP53*^mut^ categories (reds) in bulk RNA-seq LUAD tumor samples from TCGA. *P*-values were calculated using a Mann-Whitney-Wilcoxon test. **C.** Distribution of entropy scores across *TP53*^WT^ (blue), and different *TP53*^mut^ single and co-mutations (reds) in bulk RNA-seq LUAD tumor samples from TCGA. *P*-values were calculated using a Mann-Whitney-Wilcoxon test. **D.** Heatmap of mean entropy score for malignant subsets in scRNA-seq data. **E.** Stacked bar plot of proportion of HPCS cells from K and KP mice, classified into different malignant subsets from this study. **F.** Comparison of entropy scores between *KRAS*^WT^ and different *KRAS*^mut^ impact categories in A549 cells.

**Figure S5.**
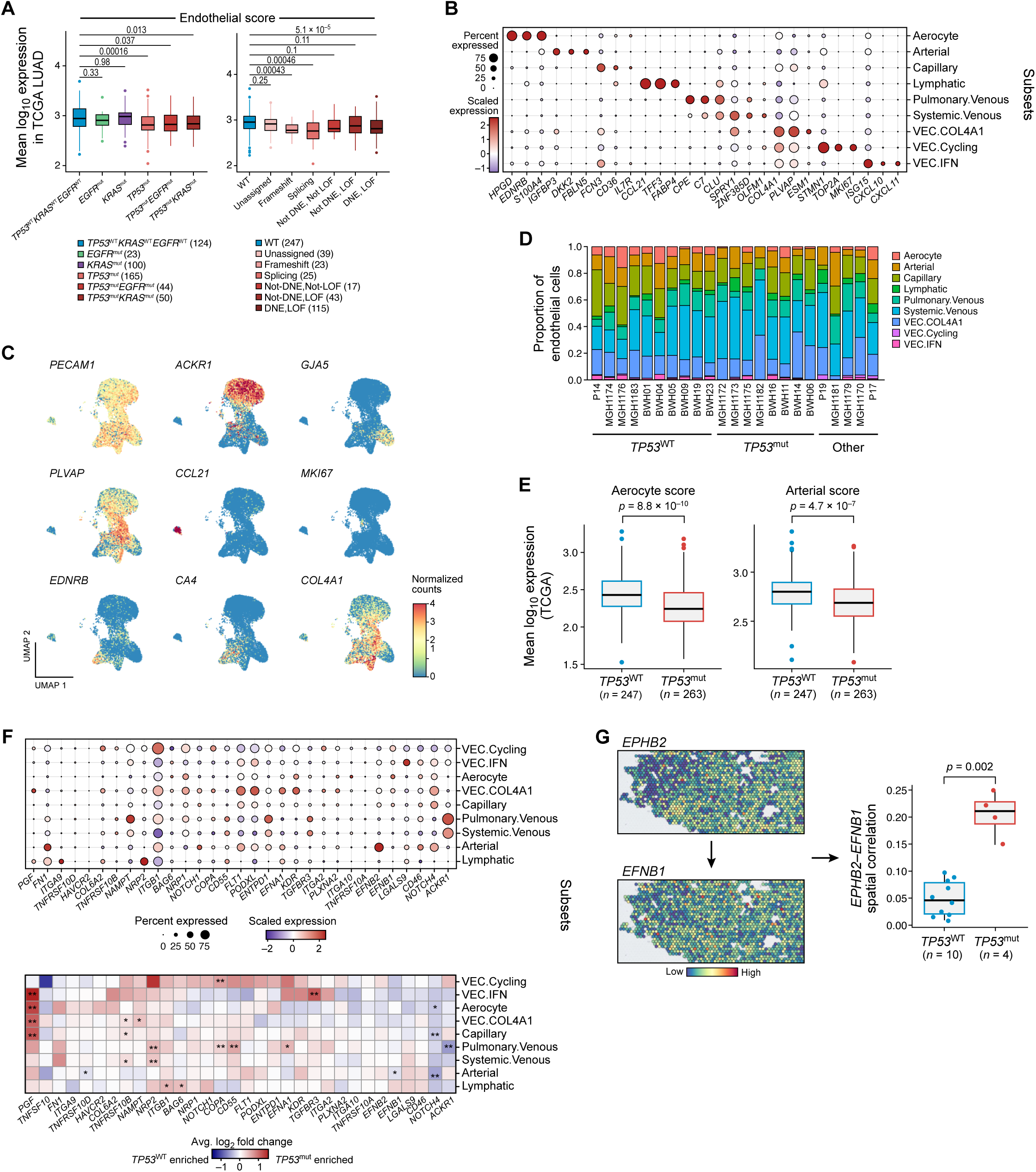
Characterizing endothelial subsets and their differential roles in *TP53*^WT^ and *TP53*^mut^ LUAD. **A.** Left: Comparison of mean log_10_ endothelial marker expression between *TP53*^WT^ and different single- or co-mutations of *EGFR*, *KRAS*, and *TP53* in bulk RNA-seq data from TCGA tumor samples. Right: Comparison of mean log_10_ endothelial marker expression between *TP53*^WT^ and different dominant-negative/loss of function *TP53*^mut^ variant categories in bulk RNA-seq LUAD tumor samples from TCGA. Mann-Whitney-Wilcoxon test was used for computing *p*-values. **B.** Dot plot showing the expression of three marker genes for each annotated endothelial subset. **C.** Feature plots of the expression of nine selected endothelial subset markers. **D.** Stacked bar plot displaying the cell composition of endothelial subsets for each scRNA-seq patient sample. **E.** Average log_10_ expression of aerocytes (left) and arterial (right) markers derived from the scRNA-seq data for *TP53*^WT^ (blue) and *TP53*^mut^ (red) LUAD bulk RNA-seq tumor samples from TCGA. *P*-values were calculated using a two-tailed multiple regression t-test. **F.** Top: Dot plot of genes involved in ligand-receptor interactions displayed in Figure 3C, Bottom: Pseudobulk differential expression analysis performed on the same genes comparing *TP53*^mut^ *vs*. *TP53*^WT^ tumor samples. Mann-Whitney-Wilcoxon test *p-*values were calculated. Red indicates enrichment in *TP53*^mut^ and blue enrichment in *TP53*^WT^ tumor samples. **G.** Left: Spatial expression of *EFHB2* and *EFNB1* in a representative *TP53*^mut^ ST sample. Right: Box plot comparing spatial correlation of *EFHB2* and *EFNB1* between *TP53*^mut^ and *TP53*^WT^ ST tumor samples. Mann-Whitney-Wilcoxon test *p*-values were computed.

**Figure S6.**
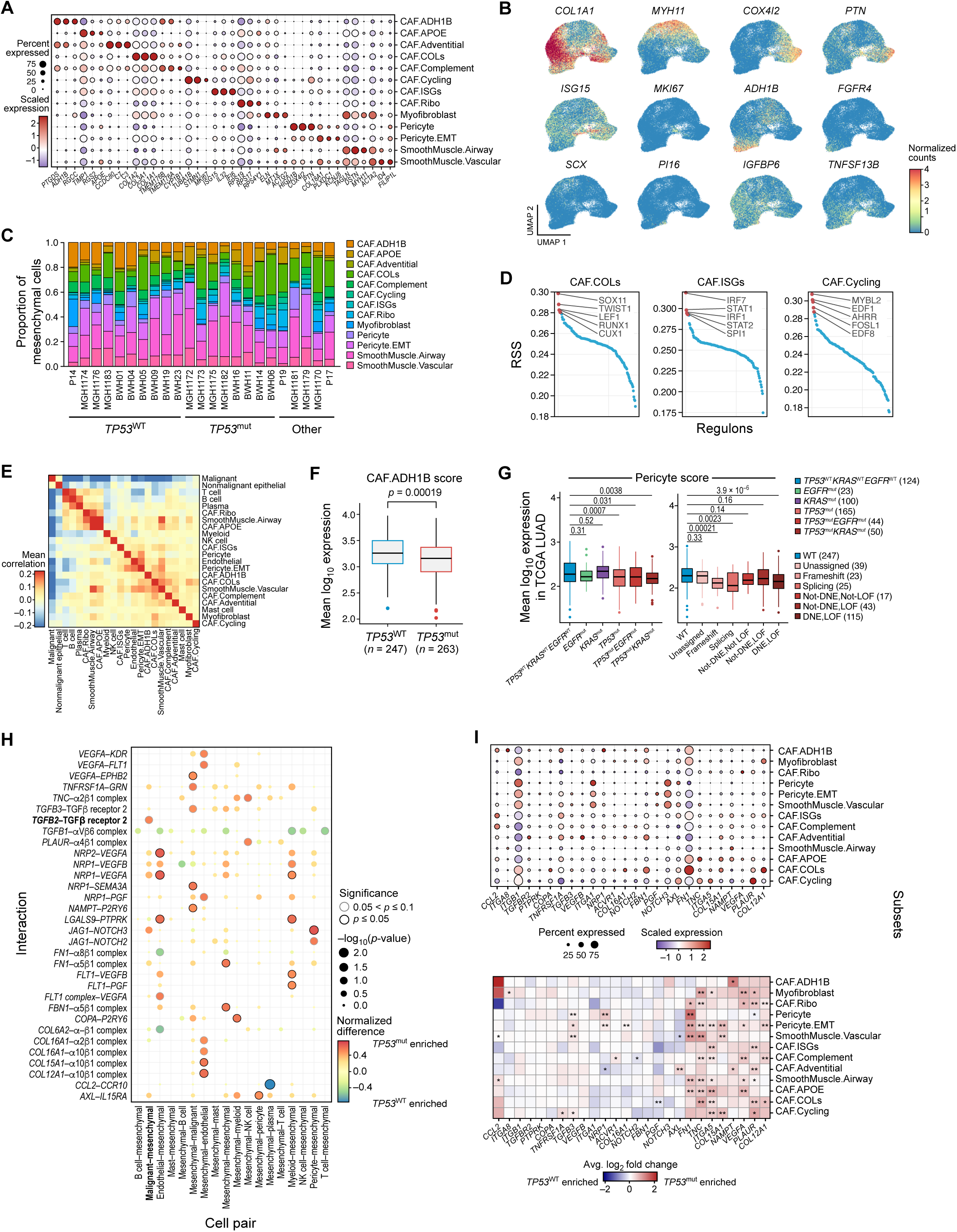
Mesenchymals cell compartment differences between *TP53*^mut^ and *TP53*^WT^ LUAD. **A.** Dot plot showing the expression of three marker genes for each annotated mesenchymal subset. **B.** Feature plot showing the expression of 12 selected mesenchymal subset markers. **C.** Stacked bar plot displaying the cell composition of mesenchymal subsets for each patient sample. **D.** Regulon specificity scores (RSS) plotted for three different mesenchymal subsets of interest. **E.** Spatial correlation (colocalization) of mesenchymal subsets with other broad cell classes across all ST tumor samples. Red indicates high colocalization and blue low colocalization between pairs of cell subsets. **F.** Average log_10_ expression of CAF.ADH1B markers derived from the scRNA-seq data for *TP53*^WT^ (blue) and *TP53*^mut^ (red) LUAD bulk RNA-seq tumor samples from TCGA. *P*-values were calculated using a two-tailed multiple regression t-test. **G.** Left: Comparison of mean log_10_ pericyte marker expression between *TP53*^WT^ and different single- or co-mutations of *EGFR*, *KRAS*, and *TP53* in bulk RNA-seq data from TCGA tumor samples. Right: Comparison of mean log_10_ pericyte marker expression between *TP53*^WT^ and different dominant-negative/loss of function *TP53*^mut^ variant categories in bulk RNA-seq LUAD tumor samples from TCGA. Mann-Whitney-Wilcoxon test was used for computing *p*-values. **H.** Dot plot of differential ligand-receptor (rows) interactions between mesenchymal cells and other cell classes (columns). Red indicates enrichment in *TP53*^mut^ and blue enrichment in *TP53*^WT^ tumor samples. Black outline indicates *p*-value ≤ 0.05, as assessed by Fisher’s exact test. **I.** Top: Dot plot of genes involved in ligand-receptor interactions displayed in **Figure S3H.** Bottom: Pseudobulk differential expression analysis performed on the same genes comparing *TP53*^mut^ *vs*. *TP53*^WT^ tumor samples. Mann-Whitney-Wilcoxon test *p-*values were calculated. Red indicates enrichment in *TP53*^mut^ and blue enrichment in *TP53*^WT^ tumor samples.

**Figure S7.**
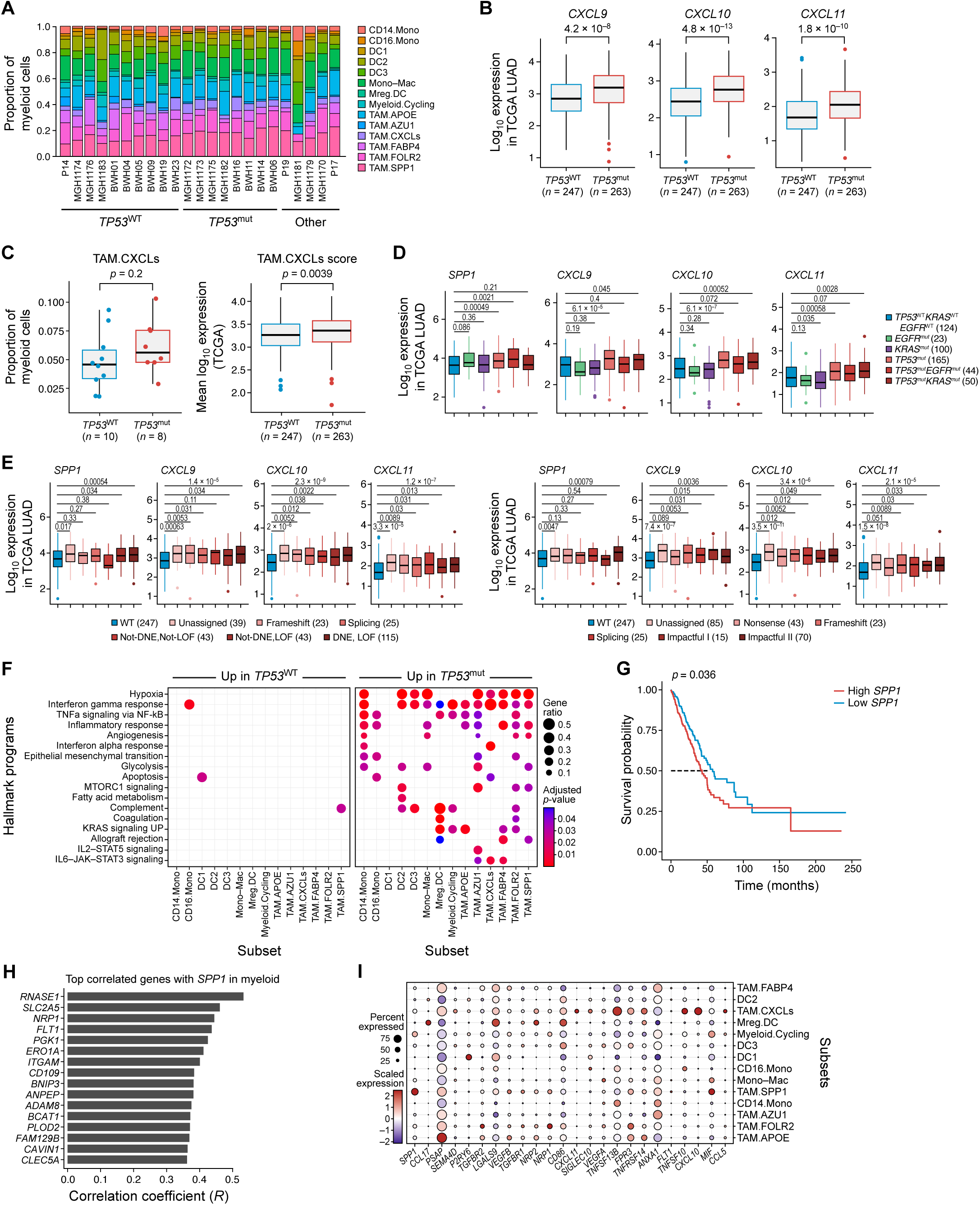
Characterization of myeloid subsets and expression of ligand-receptor genes. **A.** Stacked bar plot displaying the cell composition of myeloid subsets for each patient sample. **B.** Comparison of *CXCL9/10/11* expression in *TP53*^mut^ *vs*. *TP53*^WT^ LUAD tumor samples using bulk RNA-seq data from TCGA. **C.** Left: Proportion of TAM.CXCLs subset relative to all myeloid cells in *TP53*^WT^ (blue) *TP53*^mut^ (red) LUAD scRNA-seq tumor samples. Right: Average log_10_ expression of TAM.CXCLs markers in LUAD bulk RNA-seq tumor samples from TCGA. *P*-values were calculated using a Mann-Whitney-Wilcoxon test (scRNA-seq, left) or a two-tailed multiple regression t-test (TCGA, right). **D.** Comparison of *SPP1*, *CXCL9*/*10*/*11* gene expression among *TP53*^WT^ and different single- or co-mutations in *EGFR*, *KRAS*, and *TP53*. **E.** Comparison of gene expression among dominant-negative/loss of function (left), and impact categories (right) of *TP53*^mut^ LUAD tumor samples in TCGA. *P*-values were calculated using a Mann-Whitney-Wilcoxon test. **F.** Dot plot of gene set enrichments using the Hallmark database for the 40 most differentially expressed genes in each myeloid subsets, comparing genes up in *TP53*^WT^ (left) and up in *TP53*^mut^ (right). **G.** Kaplan-Meier survival curve showing association between *SPP1* expression and overall survival in bulk RNA-seq LUAD data from TCGA. Tumors (n = 510) were stratified into high or low expression of *SPP*1. Cox proportional-hazards model was used to calculate a *p*-value corrected for *TP53* mutational status. **H.** Horizontal bar plot showing the most correlated genes with *SPP1* expression in myeloid cells, ranked by Pearson’s correlation coefficient. **I.** Dot plot showing expression of genes involved in ligand-receptor interactions shown in Figure 4F.

**Figure S8.**
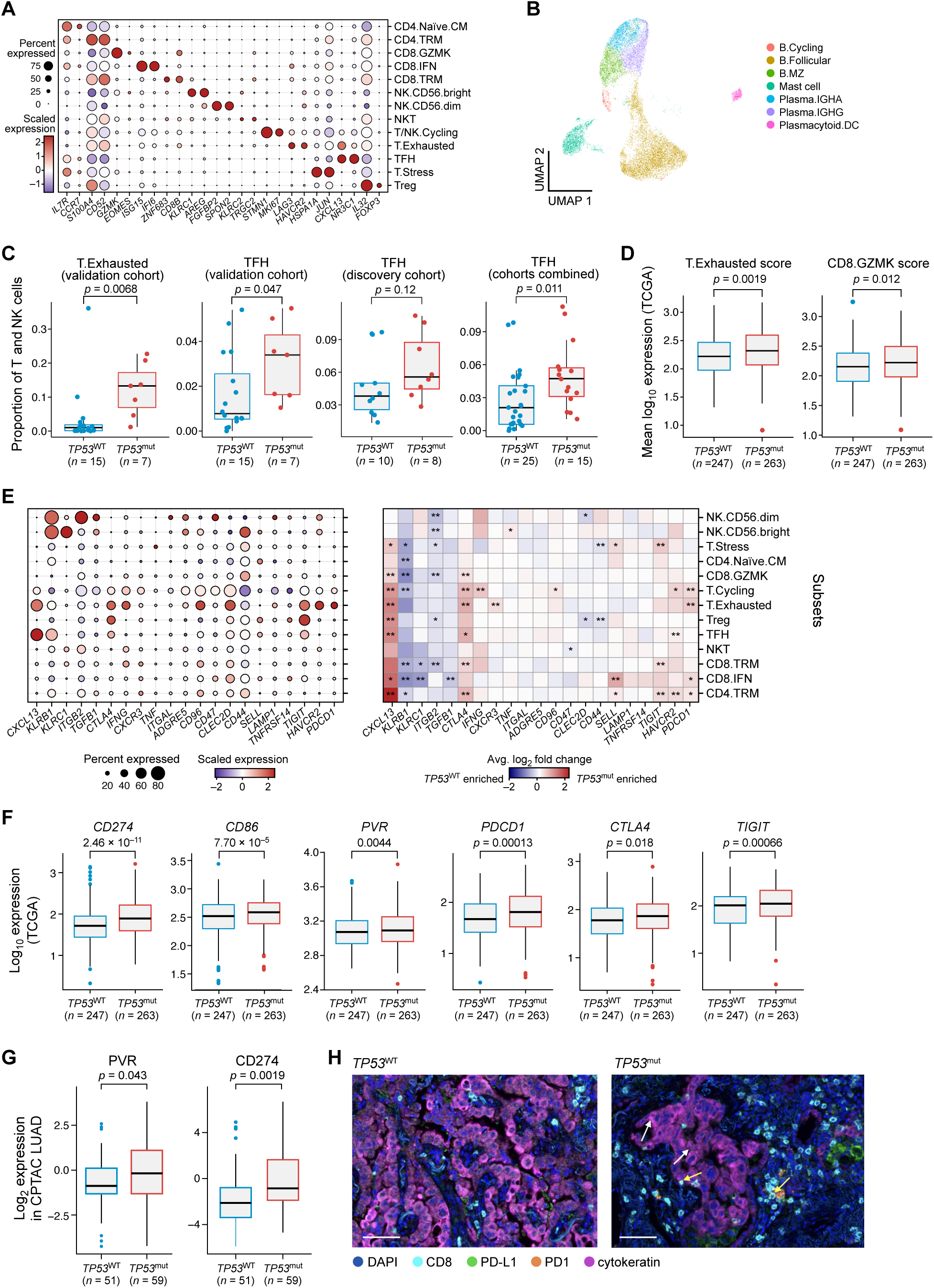
Lymphoid immune checkpoint gene expression and spatial distribution. **A.** Dot plot showing the expression of two representative marker genes for each annotated T and NK subset. **B.** UMAP of 18,520 B, plasma, mast, and plasmacytoid DC cells, integrated across tumors and colored by annotated subset. **C.** Comparison of T.Exhausted (left; validation cohort only) and TFH cell proportions (right; discovery, validation, and combined cohorts) relative to all T and NK cells between *TP53*^mut^ and *TP53*^WT^ tumors profiled by scRNA-seq. *P*-values were calculated using a Mann-Whitney-Wilcoxon test. **D.** Average log_10_ expression of specific T.Exhausted, and CD8.GZMK markers derived from the scRNA-seq data in *TP53*^WT^ (blue) and *TP53*^mut^ (red) LUAD bulk RNA-seq tumor samples from TCGA. *P*-values were calculated using a two-tailed multiple regression t-test. **E.** Left: Dot plot of genes involved in ligand-receptor analysis displayed in Figure 5D. Right: Pseudobulk differential expression analysis performed on the same genes comparing *TP53*^mut^ *vs*. *TP53*^WT^ tumor samples. Mann-Whitney-Wilcoxon test *p-*values were calculated. Red indicates enrichment in *TP53*^mut^ and blue enrichment in *TP53*^WT^ tumor samples. **F.** Log_10_ expression of *CD274*, *CD86*, *PVR*, *PDCD1, CTLA4,* and *TIGIT* in *TP53*^WT^ (blue) and *TP53*^mut^ (red) LUAD bulk RNA-seq tumor samples from TCGA. *P*-values were calculated using a two-tailed multiple regression t-test. **G.** Boxplots of PVR and CD274 (PD-L1) protein expression in CPTAC tumor samples split by *TP53* mutational status. Mann-Whitney-Wilcoxon test *p*-values were computed. **H.** Representative mIF images of additional *TP53*^WT^ (BWH04) and *TP53*^mut^ (BWH11) tumor stains. Yellow arrows correspond to PD1^+^CD8^+^ cells and white arrows correspond to PD-L1^+^cytokeratin^+^ cells. Scale bar, 50 µm.

**Figure S9.**
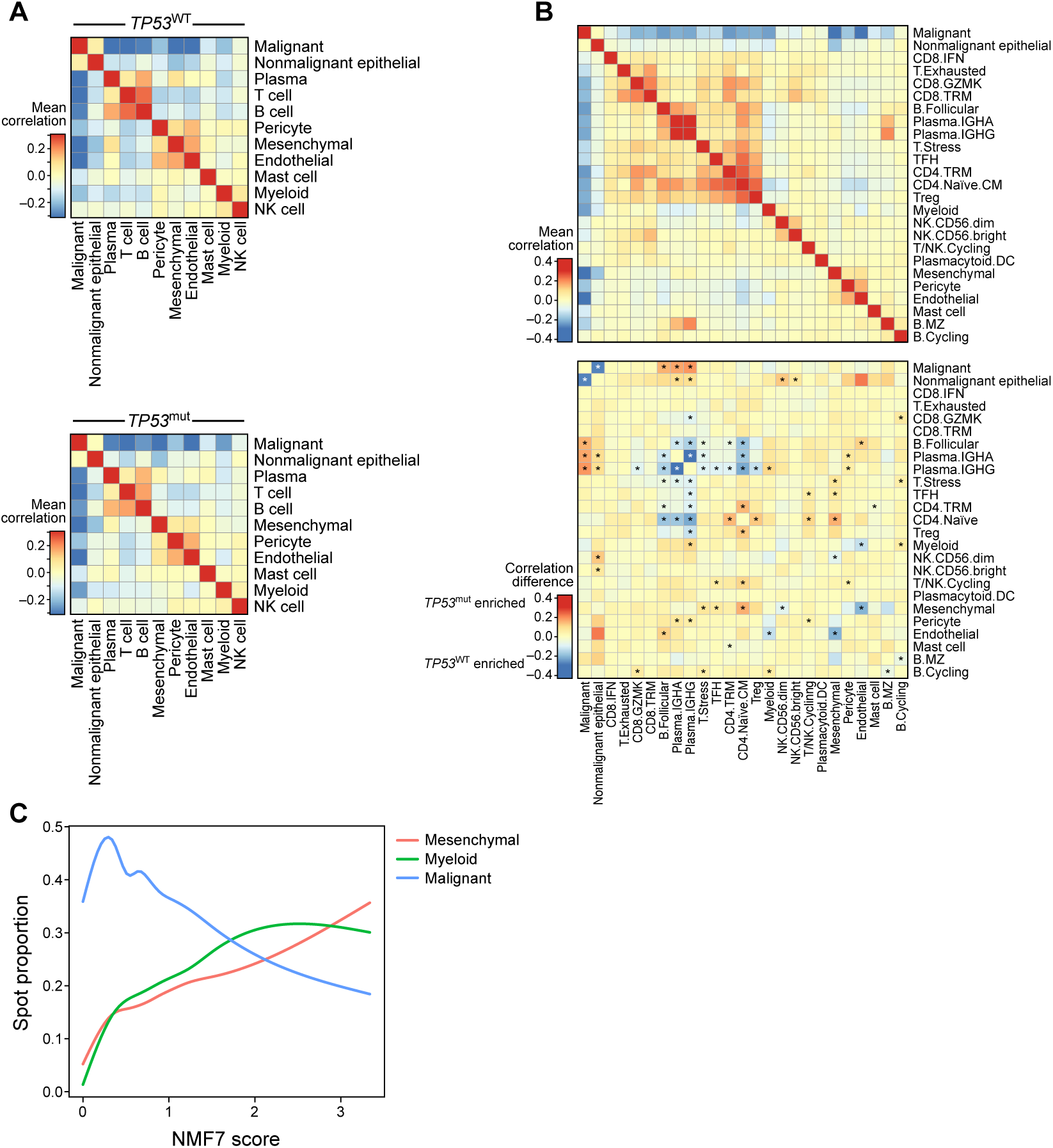
Spatial organization and cellular landmarks in *TP53*^mut^ and *TP53*^WT^ LUAD. **A.** Mean spatial correlation of broad cell classes across ST spots averaged across *TP53*^WT^ (top) and *TP53*^mut^ (bottom) samples. **B.** Mean (top) and differential (bottom) spatial correlation heatmap of lymphoid (B, plasma, T, NK) subsets with other broad cell classes between *TP53*^mut^ and *TP53*^WT^. Red indicates either high overall colocalization (top) or colocalization enrichment in *TP53*^mut^ (bottom) while blue represents either low overall colocalization (top) or colocalization enrichment in *TP53*^WT^ (bottom) tumor samples between pairs of cell subsets. **C.** Line plot showing proportion of malignant, myeloid, and mesenchymal cells ordered from NMF7 low spots to NMF7 high spots (x-axis).

**Table S1. Clinical data for patient cohort**

**Table S2. High level cell class markers and subset markers**

**Table S3. Clinical data for validation cohort**

